# Somatic evolution in non-neoplastic IBD-affected colon

**DOI:** 10.1101/832014

**Authors:** Sigurgeir Olafsson, Rebecca E. McIntyre, Tim Coorens, Timothy Butler, Hyunchul Jung, Philip Robinson, Henry Lee-Six, Mathijs A. Sanders, Kenneth Arestang, Claire Dawson, Monika Tripathi, Konstantina Strongili, Yvette Hooks, Michael R. Stratton, Miles Parkes, Inigo Martincorena, Tim Raine, Peter J. Campbell, Carl A. Anderson

## Abstract

Inflammatory bowel disease (IBD) is a chronic inflammatory disease associated with increased risk of gastrointestinal cancers. Here, we whole-genome sequenced 447 colonic crypts from 46 IBD patients, and compared these to 412 crypts from 41 non-IBD controls. The average mutation rate of affected colonic epithelial cells is 2.4-fold that of healthy colon and this increase is mostly driven by acceleration of mutational processes ubiquitously observed in normal colon. In contrast to the normal colon, where clonal expansions outside the confines of the crypt are rare, we observed widespread millimeter-scale clonal expansions. We discovered non-synonymous mutations in *ARID1A, FBXW7, PIGR and ZC3H12A,* and genes in the interleukin 17 and Toll-like receptor pathways, under positive selection in IBD. These results suggest distinct selection mechanisms in the colitis-affected colon and that somatic mutations potentially play a causal role in IBD pathogenesis.

## Introduction

Inflammatory bowel disease (IBD) is a debilitating disease characterized by repeated flares of intestinal inflammation. The two major subtypes of IBD, Crohn’s disease (CD) and ulcerative colitis (UC), are distinguished by the location, continuity and nature of the inflammatory lesions. UC affects only the large intestine, spreading continuously from the distal to proximal colon, whereas CD most commonly affects the small and large intestine, and is characterized by discontinuous patches of inflammation. In addition to the significant morbidity associated with the disease, IBD patients have a 1.7-fold increased risk of developing gastrointestinal cancers compared to the general population. Cancer risk is associated with the duration, extent and severity of disease and cancers tend to occur earlier in life in IBD patients (Lutgens *et al*., 2013; Beaugerie and Itzkowitz, 2015; Adami *et al*., 2016). As a result, patients require regular endoscopic screening and may undergo prophylactic colectomy to mitigate this risk (Beaugerie and Itzkowitz, 2015; Adami *et al*., 2016).

That somatic mutations contribute to the development of cancer is well established, but their patterns, burden and functional consequences in diseases other than cancer have not been extensively studied. Methodological developments have now enabled the analysis of polyclonal somatic tissues, allowing characterization of somatic mutations in normal tissues such as skin (Martincorena *et al*., 2015), esophagus (Martincorena *et al*., 2018; Yokoyama *et al*., 2019), endometrium (Moore *et al*., 2018; Suda *et al*., 2018), lung (Yoshida *et al*., 2020) and colon (Blokzijl *et al*., 2016; Lee-Six *et al*., 2019). In the setting of non-neoplastic diseases, chronic liver disease has had the most attention, with studies showing that compared to healthy liver, hepatic cirrhosis is associated with acquisition of new mutational processes, increased mutation burden and larger clonal expansions (Brunner *et al*., 2019; Kim *et al*., 2019; Zhu *et al*., 2019).

Colonic epithelium is well suited to the study of somatic mutations on account of its clonal structure. It is organized into millions of colonic crypts, finger-like invaginations composed of approximately 2000 cells (Potten *et al*., 1992) each extending into the lamina propria below.

At the base of each crypt reside a small number of stem cells undergoing continuous self-renewal through stochastic cell divisions (Lopez-Garcia *et al*., 2010; Snippert *et al*., 2010). As a result, the progeny of a single stem cell iteratively sweeps through the entire niche and the epithelial cells that line the crypt are the progeny of this single clone. Active IBD disrupts these normal stem cell dynamics - the epithelial lining is damaged, the organized crypt structure is ablated and the barrier between lumen and mucosa is disrupted.

We hypothesized that the recurrent cycles of inflammation, ulceration and regeneration seen in inflammatory bowel disease could impact the mutational and clonal structure of intestinal epithelial cells. To test these hypotheses we isolated and whole-genome sequenced 447 colonic crypts from 46 IBD patients with varying degrees of colonic inflammation, both active and previous, and compared the mutation burden, clonal structure, mutagen exposure and driver mutation landscape to colonic crypts from healthy donors (Lee-Six *et al*., 2019).

## Results

### IBD more than doubles the mutation rate of normal colonic epithelium

We used laser capture microdissection (LCM) to isolate 447 colonic crypts from endoscopic biopsies taken from 28 UC patients and 18 CD patients (Supplementary Table 1 and Supplementary Figure 1). Biopsies were annotated as never-, previously- or actively inflamed at the time of sampling (Methods). The dissected crypts were whole-genome sequenced to a median depth of 18.2X, allowing us to call somatic substitutions, small insertions and deletions (indels), larger copy number changes, somatic retrotranspositions and aneuploidies affecting whole chromosomes or chromosome arms (Methods, Supplementary Tables 2 through 6).

To assess if IBD is associated with a difference in the mutation burden of the colonic epithelium, we combined our data with data from 412 crypts sequenced as part of our recent study of somatic mutations in normal colon (Lee-Six *et al*., 2019) (hereafter referred to as the control data). We fitted linear mixed-effects models (LMMs) to estimate the independent effects of age, disease duration and biopsy location on mutation burden, while controlling for the within-patient and within-biopsy correlations inherent in our sampling strategy (Supplementary Table 3, Methods). We estimated the effect of IBD to be 54 substitutions per crypt per year of disease duration (33-74 95% CI, P=6.7×10^-7^, LMMs and likelihood ratio test - Figure 1). These mutations are in addition to the 40 (31-50, 95% CI) substitutions we estimated are accumulated on average per year of life under normal conditions, suggesting that mutation rates are increased ∼2.3-fold in regions of the IBD-affected colon on average. Compared to controls, patients with IBD had greater between-patient variance in mutation burden (SD=900 versus 112 mutations for cases and controls, respectively. P=5.8×10^-8^, LMMs and likelihood ratio test) and greater within-patient variance (SD = 1067 versus 390 mutations for cases and controls, respectively. P=0.015). The increased between-patient variance likely reflects differences in disease severity and response to treatment among patients; the increased within-patient variance probably reflects region-to-region differences in disease severity along the colon. We similarly estimated an increase in the indel burden in IBD, with an excess of seven indels per crypt per year of IBD (5 - 9 95% CI, P=1.0×10^-10^ - Figure 1) in addition to the estimated 1 (0.3-1.7 95% CI) indel that is accumulated per crypt per year of life. We found no significant difference in the mutation burden between UC and CD patients.

**Figure 1.**
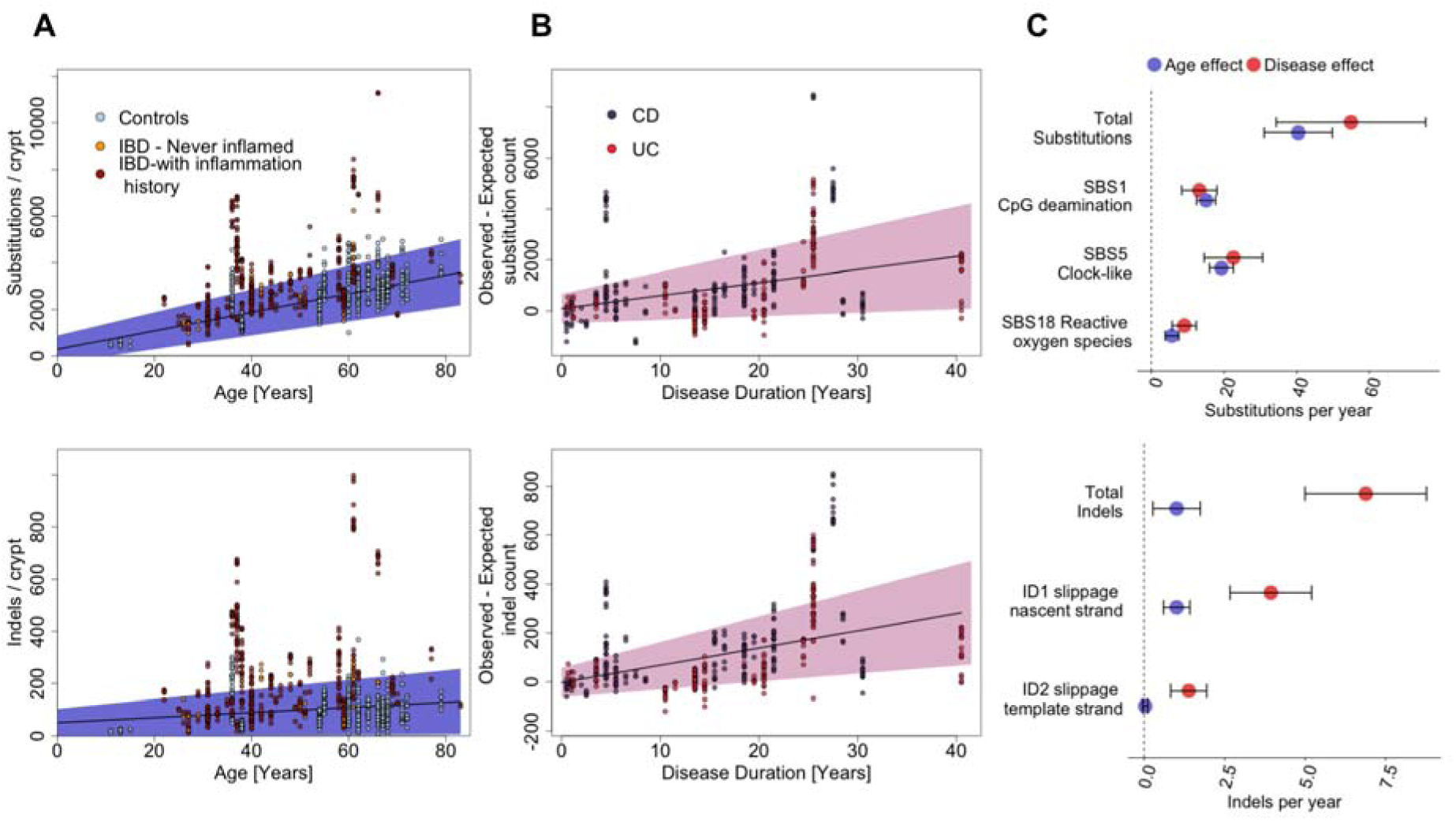
Mutation burden in the IBD colon. **A**) Substitution (top) and indel (bottom) burden as a function of age. Each point represents a colonic crypt and is coloured by disease status. The line shows the effect of age on mutation burden as estimated by fitting a linear mixed effects model, correcting for sampling location, sequencing coverage and the within-biopsy and within-patient correlation structure, considering both IBD cases and controls. The blue shaded area represents the 95% confidence interval of the age effect estimate. **B**) Estimated excess of substitutions (top) and indels (bottom) in crypts from IBD patients as function of disease duration. **C**) A comparison of the effects of age and disease duration on the total mutation burden and on the burden of mutational signatures that associate with IBD duration. Error bars represent the 95% confidence intervals of the estimates. IBD: Inflammatory bowel disease. CD: Crohn’s disease. UC: Ulcerative colitis. SBS: Single base substitution signature. ID: Indel signature.

Smoking status was available for a subset of the IBD cohort (362 crypts from 35 patients). In this restricted dataset, we found an effect of smoking duration of 49 (16 - 84 95% CI, P=0.0037) substitutions per crypt per year of smoking. The effect of disease duration was unchanged, suggesting this effect is not driven by differences in smoking habits between cases and controls. Smoking has been reported to increase the risk of CD and be protective for UC (Mahid *et al*., 2006) but we found no interaction effect between smoking and disease type (P=0.63). Smoking status was not available for the control cohort.

### IBD accelerates age-related mutational processes

The somatic mutations found in the cells of a colonic crypt reflect the mutational processes that have acted on the stem cells and their progenitors since conception. Distinct mutational processes each leave a characteristic pattern, a mutational signature, within the genome, distinguished by the specific base changes and their local sequence context (Alexandrov *et al*., 2013, 2020). We extracted mutational signatures jointly for IBD and control crypts and discovered 12 substitution signatures (SBS) and five indel signatures (ID), all of which have been previously observed in tissues from individuals without IBD (Methods, Supplementary Table 2, Supplementary Figures 3 and 4). Comparing our IBD cases and controls, we found that approximately 80% of the increase in mutation burden in cases is explained by signatures that are also found ubiquitously in normal colon (Blokzijl *et al*., 2016; Lee-Six *et al*., 2019). These are substitution signatures 1, 5 and 18 and indel signatures 1 and 2, as defined by Alexandrov et al. (Figure 1), which cause an increase of 13 (8-18 95% CI), 23 (15-31 95% CI) and 9 (6-12 95% CI) substitutions per crypt per year of disease, respectively (P=3.3×10^-7^, 2.1×10^-7^ and 4.5×10^-7)^, and 4.4 (3.3-5.5 95% CI) and 1.8 (1,2-2.4, 95% CI) indels per crypt per year, respectively (P=8.4 ×10^-12^ and P=1.4 ×10^-7^, LMMs and likelihood ratio tests). Substitution signatures 1 and 5 are clock-like and thought to be associated with cell proliferation, while signature 18 has been linked with reactive oxygen species (Alexandrov *et al*., 2020). The indel signatures ID1 and ID2 are both thought to be the result of polymerase slippage during DNA replication (Alexandrov *et al*., 2020).

The remaining 20% of the increase in substitution burden is a consequence of rarer mutational processes and treatment. For example, we found 96 crypts with over 150 mutations attributed to purine treatment in a subset of seven IBD patients, five of whom have a documented history of such treatment. However, the number of mutations attributed to purine was not associated with purine therapy duration, and some patients showed large mutation burdens despite brief, or indeed no, documented exposure. For example, one patient received azathioprine for two weeks and mercaptopurine for two weeks and had significant adverse reactions to both drugs. This brief treatment resulted in a median of 204 mutations (range: 120-374) attributed to purine treatment in the crypts from this individual. Other patients had long term exposure to azathioprine without accruing any purine-related mutations (Figure 2B). There was also great within-patient variation in the burden of the purine signature. The largest range was observed for patient 40, which has a 7 year history of purine treatment. The estimated burden of the purine signature in crypts from this patient ranged from 69 to 1005.

**Figure 2:**
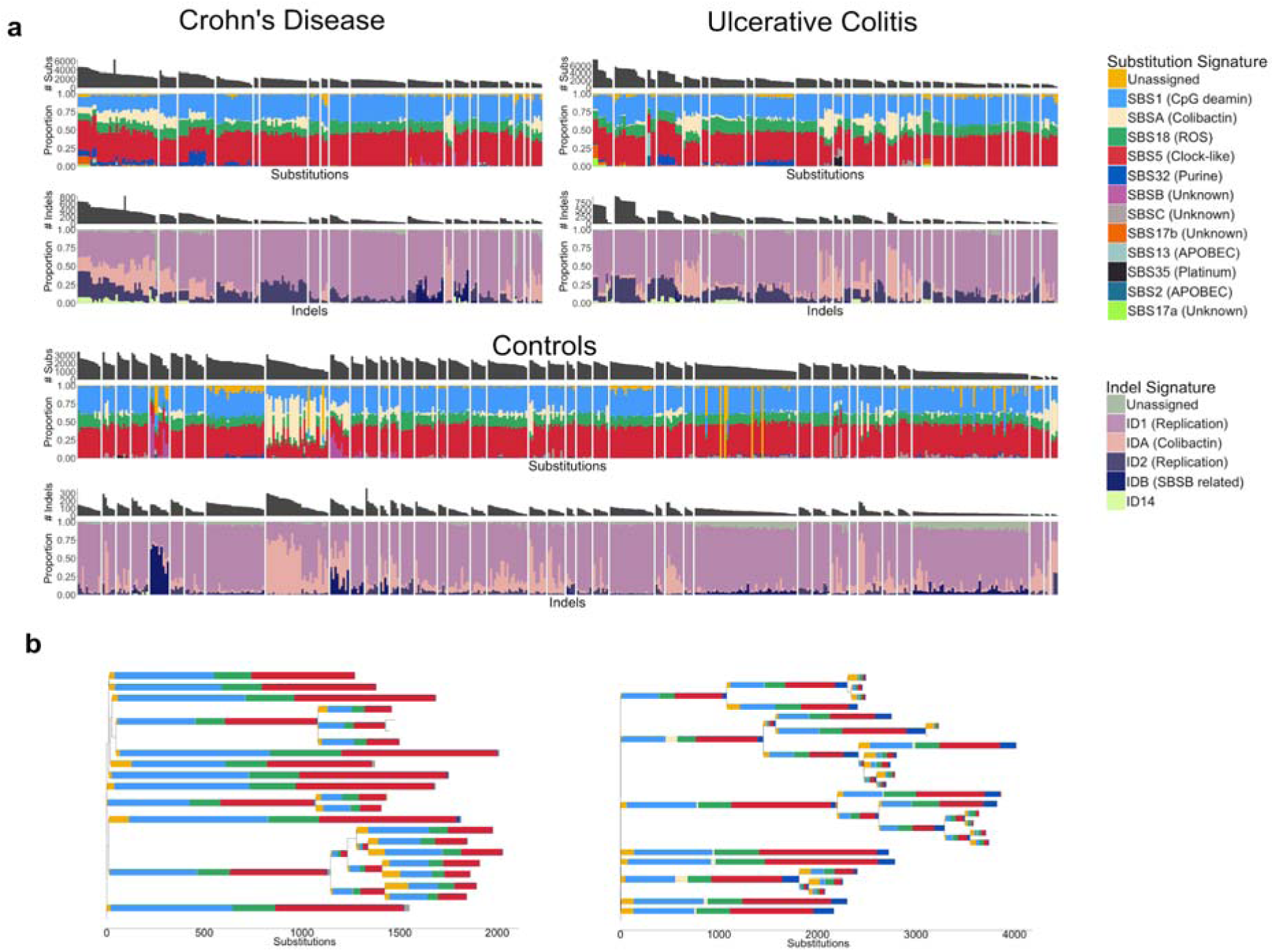
Mutational signatures in colonic crypts. a) A stacked barplot showing the proportional contribution of single-base-substitution (SBS) signatures (Top) and Indel (ID) signatures (Bottom) to the mutation burden of each crypt. Crypts are grouped by patient and crypts from CD, UC and controls are shown separately. Signature nomenclature is the same as in Alexandrov et al (2020). The ‘Unassigned’ component represents uncertainty of the signature extraction. b) Phylogenetic trees of two patients with widespread ulcerative colitis. The colours of the branches reflect the relative contribution of each mutational signature extracted for those branches as in a). The patient on the left has received azathioprine treatment for 10 years but shows no SBS32 burden (dark blue). In contrast, the patient on the right received azathioprine for 2 weeks and mercaptopurine for 2 weeks and had significant adverse reactions to both drugs. SBS32 is found in most crypts from this patient. All crypts are from inflamed biopsies.

Five signatures previously discovered in the normal colon (Lee-Six *et al*., 2019), SBSA, SBSB and SBSC, IDA and IDB were also present in the context of IBD. SBSA and IDA and SBSB and IDB are highly correlated (Supplementary Figure 4) and likely represent the same underlying mutational processes. SBSA and IDA are of particular interest since they have recently been shown to be caused by the genotoxin colibactin, which is produced by bacteria harboring a polyketide synthases (pks) pathogenicity island (Pleguezuelos-Manzano *et al*., 2020). *pks*+ *E. coli* have been reported at increased frequency in IBD (Arthur *et al*., 2012), but we found no relationship between SBSA or IDA burden and disease status or disease duration after correcting for higher burden of both in the left-side of the colon (the site primarily affected in UC). As in normal colon, SBSA and SBSB were primarily found in early branches of the phylogenetic trees (Supplementary Figures 5 and 6). Signatures SBSB, SBSC and SBS32 have not been reported in studies of sporadic colorectal cancers (Alexandrov *et al*., 2020), perhaps due to the comparative complexity and diversity of cancer mutation profiles. SBS32 however, would only be expected in patients receiving purine therapy and so would not be present in sporadic colorectal cancers. These signatures have also not been reported in studies of colitis-associated colorectal cancers but this is likely due to a relative lack of power due to the small number of sequenced exomes (Robles *et al*., 2016; Baker *et al*., 2018; Din *et al*., 2018).

Signatures 2 and 13, which are associated with APOBEC activity, and signatures 17a and 17b, which are of unknown aetiology, were active in a small number of crypts with high mutation burdens. SBSB, SBSC, SBS17a/b and SBS2/SBS13 are too rare for us to be powered to detect any difference between IBD and controls or to associate these with any clinical feature documented in our metadata. Finally, we found signature 35, associated with platinum compound therapy, in one patient with a history of platinum treatment for squamous cell carcinoma of the tongue. The patient received 40mg/m^2^ of cisplatin therapy on a weekly basis. He completed three of six planned treatment cycles with therapy termination due to toxicity. This relatively brief treatment resulted in a medium of 430 mutations (range 350-461) per crypt that were attributed to signature 35, equivalent to about 10 years of normal mutagenesis.

### IBD associates with the burden of structural variants

We called copy number variants (CNVs), somatic retrotranspositions and loss-of-heterozygosity events affecting whole chromosomes or chromosome arms (referred to here as aneuploidies for simplicity) in both our IBD cases and non-IBD controls. The burden of structural variants is modest in both datasets (Figure 3) but for IBD, we identified the occasional clone that carried a large number of CNVs and retrotranspositions (Figure 3a and b). The numbers of CNVs and retrotranspositions are associated with IBD duration. We estimated the CNV mutation rate to be 0.67 CNVs per crypt per year of disease (0.027 - 0.11 95% CI, P=9.4 x 10^-4^, Likelihood ratio test of mixed-effects Poisson regressions) and the retrotransposition mutation rate to be 0.061 (0.014 - 0.11, 95% CI, P=0.012). This corresponds to one CNV per crypt every 15 years of disease duration and one retrotransposition event every 16.4 years of disease duration on average. However, a handful of clones accumulated many structural variants (SVs), while the majority had none, suggesting that the processes driving their acquisition may be episodic rather than continuous. This would be in line with findings from other reports linking rapid accrual of SVs with the transition from normal to dysplastic mucosa (Baker *et al*., 2018) and cancers accruing copy number gains in a punctuated manner (Gerstung *et al*., 2020).

**Figure 3:**
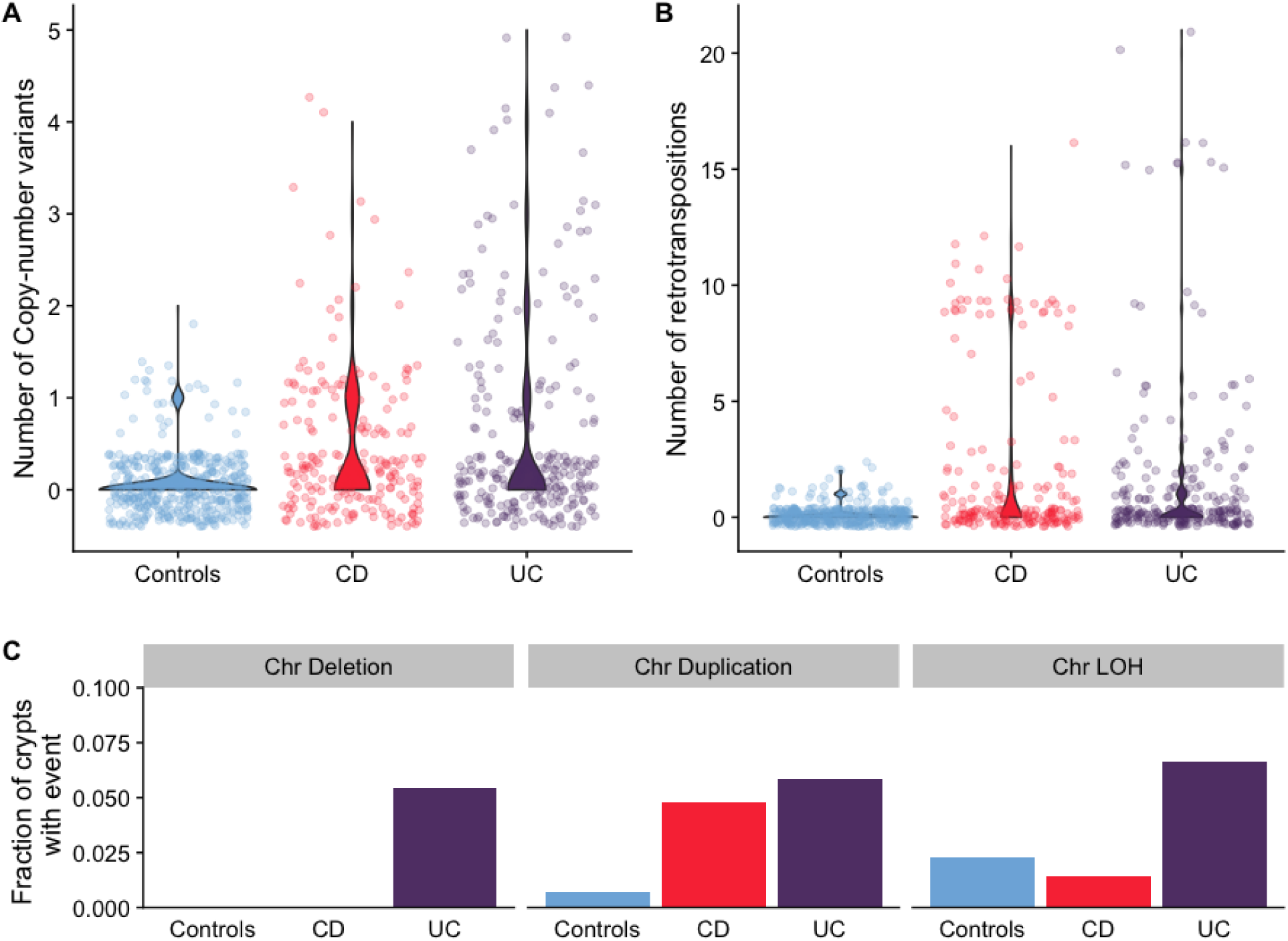
Burden of structural variants in inflammatory bowel disease affected colon compared with IBD-unaffected colon. a) Number of copy number variants in IBD sub-types compared with controls. b) Number of somatic retrotranspositions in IBD sub-types compared with controls. c) Fraction of crypts with inflammation history that carry chromosomal aneuploidies.

We found a higher fraction of IBD crypts carrying aneuploidies than in controls (43/419 compared with 13/412, Figure 2c). However, this was driven by large clones carrying aneuploidies and the number of events was not significantly associated with disease duration (P=0.38). The number of retrotranspositions and aneuploidies is associated with higher substitution burden (49 (27-72 95% CI), P = 2.1 x 10^-5^ and 207 (75-338 95% CI), P = 2.1 x 10^-3^, respectively) and retrotranspositions and CNVs are associated with higher indel burden (9 (3-12 95% CI), P = 2.3 x 10^-9^ and 12 (4-19, 95% CI), P = 2.5×10^-3^, respectively).

### IBD creates a patchwork of millimeter-scale clones

Colonic crypts divide by a process called crypt fission, whereby a crypt bifurcates at the base and branching elongates in a zip-like manner towards the lumen. This process is relatively rare in the normal colon, wherein each crypt fissions on average only once every 27 years (Nicholson *et al*., 2018; Lee-Six *et al*., 2019). Compared to normal colon, we found much larger clonal expansions in IBD patients, evident of numerous crypt fission events occurring late in molecular time. We observed several examples of individual clones spanning entire 2-3 mm endoscopic biopsies (Figure 3, Figure 4a, Supplementary Figures 5 and 6). Our ability to estimate clone sizes is restricted by the small size of the biopsies. However, when we biopsied the same inflamed or previously inflamed region more than once, on only one occasion out of 19 biopsy pairs did we observe a clone stretching between biopsies that were taken a few millimeters apart (Supplementary Figures 5 and 6), while most biopsies contained more than one clone. To improve our ability to detect larger clones, we sampled three patients more broadly. Nine biopsies, forming a 3×3 grid with 1 cm separating biopsies, were obtained from each patient. We dissected 187 crypts from the biopsies and performed whole exome sequencing on individual crypts. Phylogenetic trees were reconstructed based on somatic mutations identified (Figure 4b-d). While clonal expansions within biopsies are common, we found clones extending between neighboring biopsies in only one of these patients. A substantial body of evidence exists documenting widespread clonal expansions giving rise to dysplasia and ultimately to colorectal cancer in IBD (reviewed in (Choi *et al*., 2017)). Colitis-associated colorectal cancers, which are enriched with synchronous lesions (Lam, Chan and Leung, 2014; Choi *et al*., 2015), commonly grow from a background of a pre-cancerous field which has expanded many centimeters or even the whole length of the colon (Leedham *et al*., 2009; Galandiuk *et al*., 2012). Mutations in *TP53* are thought to be especially prominent in the growth of these clones but aneuploidies and *KRAS* mutations are also commonly observed (Holzmann *et al*., 1998; Leedham *et al*., 2009; Galandiuk *et al*., 2012). In our material of non-dysplastic tissue from individuals without colorectal neoplasia, we find smaller clones and mutations in *TP53*, *KRAS* or *APC* are rare. In summary, IBD-affected regions are generally not dominated by a single major clone, but are more accurately viewed as an oligoclonal patchwork of clones that often grow considerably larger than in healthy colon.

**Figure 4.**
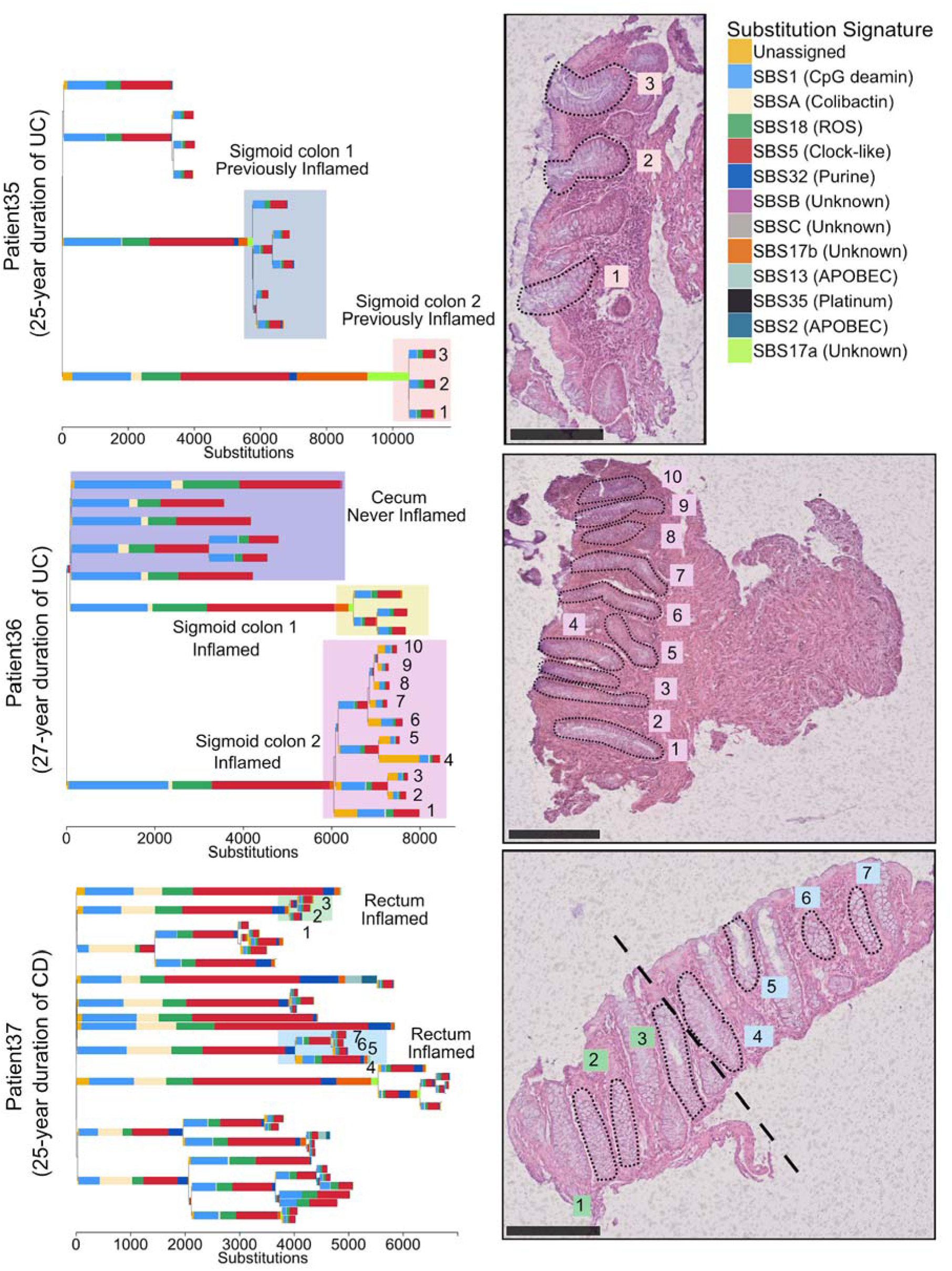
Examples of clonal expansions in three IBD patients. (Top) A phylogenetic tree of crypts sampled from a 66 year old patient with a 25 year history of ulcerative colitis. The accompanying biopsy image shows the crypts from the orange shaded area. The clones highlighted in blue and orange come from the same previously inflamed site and were millimeters apart. A large difference in the mutation burden of these clones is driven by a local activation of signatures 17a and 17b in the orange shaded clone. (Middle) A phylogenetic tree of crypts sampled from a 61 year old patient with a 27 year history of ulcerative colitis. The clones highlighted in purple and yellow come from biopsies taken millimeters apart. The accompanying biopsy image shows the crypts from the purple clone. (Bottom) A phylogenetic tree of crypts sampled from a 37 year old patient with a 25 year history of Crohn’s disease affecting the colon. A biopsy overlaps two clones (in blue and green).

### Distinct patterns of selection in IBD compared with normal epithelium

The recurrent cycles of inflammation and remission which characterise IBD could create an environment in which clones containing advantageous mutations may selectively spread in the mucosa. This advantage may manifest either through faster cell division and elevated crypt fission rate or through increased resistance to the cytotoxic effects of inflammation. To identify mutations which likely confer selective advantage on the cell, we searched for mutations occurring in canonical mutation hotspots from the Cancer Genome Atlas (Supplementary Table 7). This revealed a total of 10 missense mutations in *KRAS*, *BRAF, TP53*, *ERBB2, ERBB3* and *FBXW7* occurring at canonical hotspots (Supplementary Table 8). Additionally, we found a heterozygous nonsense mutation in *APC* and frameshift indels in known colorectal tumour suppressors; *ATM, SOX9, RNF43* and *ZFP36L2,* of likely driver status (Figure 4A and Supplementary Table 8). Furthermore, two large-scale deletions in our dataset overlap known tumour suppressors, *PIK3R1* and *CUX1*, and are likely drivers (Supplementary Figure 7). The number of putative cancer drivers found in a crypt is associated with increased burden of both substitutions (342 substitutions per driver, 173-512 95% CI, P=1.4 x 10^-4^) and indels (52 indels per driver, 30-75 95% CI, P=2.1 x 10^-5^), as well as with each of the replication-related signatures (SBS1, SBS5, SBS18, ID1 and ID2, Supplementary Table 9). The strongest association however, was with the purine signature (SBS32). We estimated the burden of purine signature to be increased by 94 (73-116, 95% CI, P = 2.9 x 10^-16^) substitution per driver, suggesting that rapidly dividing cells may be particularly susceptible to the mutagenic effect of purine treatment.

To search for genes under positive selection, we assessed the ratio of non-synonymous to synonymous mutations (dN/dS) across all IBD crypts, while correcting for regional and context-dependent variation in mutation rates (Martincorena *et al*., 2017). Genes with dN/dS ratios significantly different from 1 are considered to be under selective pressure. This analysis revealed four genes, *ARID1A, FBXW7, PIGR* and *ZC3H12A*, to be under positive selection in the IBD colon (Figure 6A and Supplementary Figure 8). *ARID1A* and *FBXW7* are well-established tumor suppressors and are found mutated at similar frequencies in sporadic- and colitis-associated colorectal cancers (Martincorena *et al*., 2017; Baker *et al*., 2018). In several instances, distinct heterozygous mutations in the same gene were found in different crypts from the same patient (Supplementary Figures 4 and 5). For example, in one patient suffering from pan-colitis we found four distinct *PIGR* mutations in four biopsies from the right, transverse and left side of the colon (Figure 6B). We did not detect a significant signal of selection of mutation in the two genes, *AXIN2* or *STAG2,* which we previously found to be under positive selection in the normal colon (Lee-Six *et al*., 2019) (P = 0.98 and 0.74, respectively) nor was there any evidence of selection of *PIGR* or *ZC3H12A* mutants in the normal colon (Supplementary Table 10). We did not find a significant difference in the mutation burden of any of these genes between UC and CD, suggesting that similar selection pressures are operative in mucosal tissue in both diseases.

**Figure 5:**
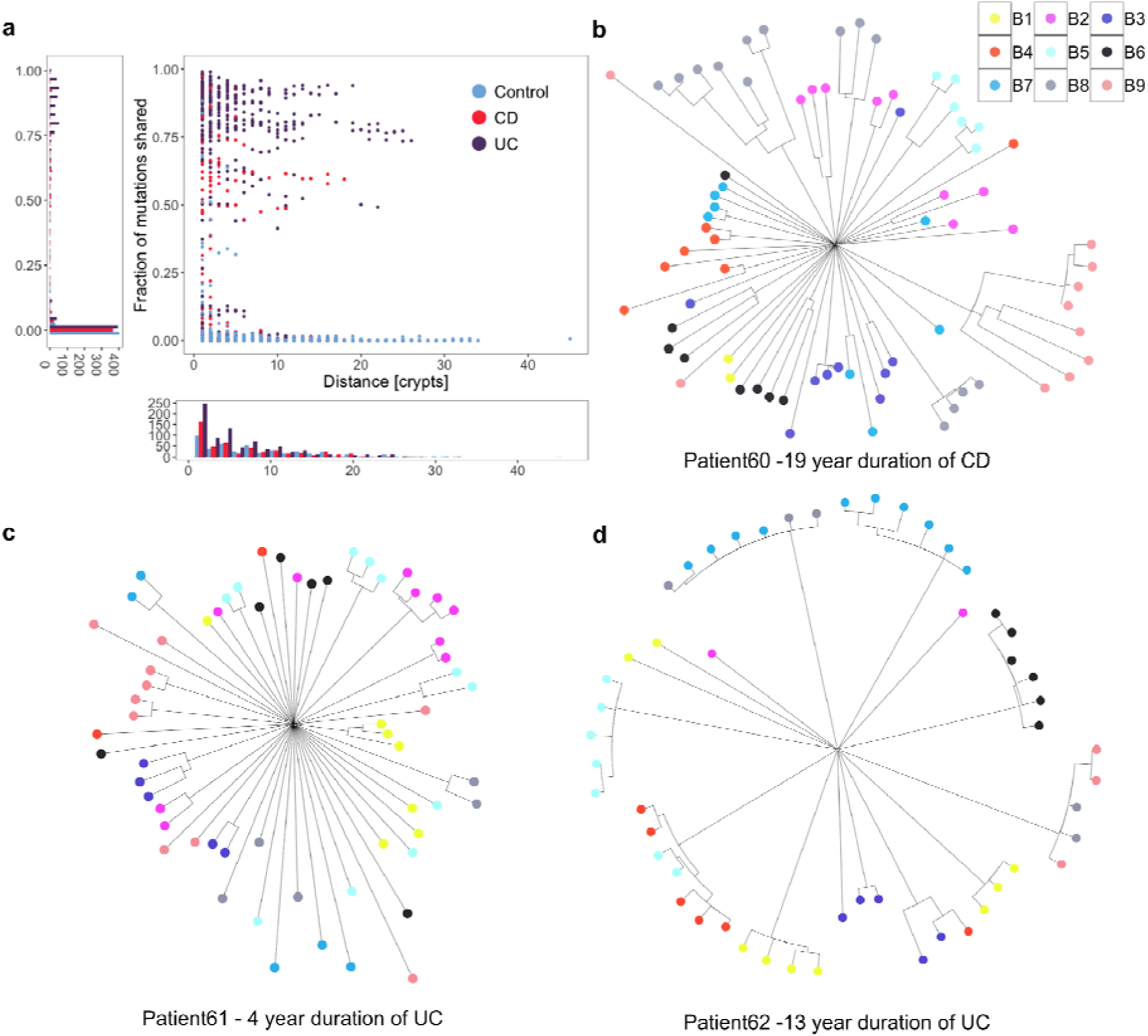
Clonal structure of the IBD colon. a) For pairs of crypts from the same biopsy, the figure shows the number of mutations that are shared between a pair as a fraction of the average mutation burden of the two crypts and this is plotted as a function of the distance between the pair. b) A phylogenetic tree showing crypts sampled from 9 biopsies from the sigmoid colon of a 36 year old male diagnosed with CD 19 years prior to sampling. c) A phylogenetic tree showing crypts sampled from 9 biopsies from the rectum of a 71 year old male diagnosed with UC 4 years prior to sampling. d) A phylogenetic tree showing crypts sampled from 9 biopsies from the rectum of a 42 year old female diagnosed with UC 13 years prior to sampling.

**Figure 6.**
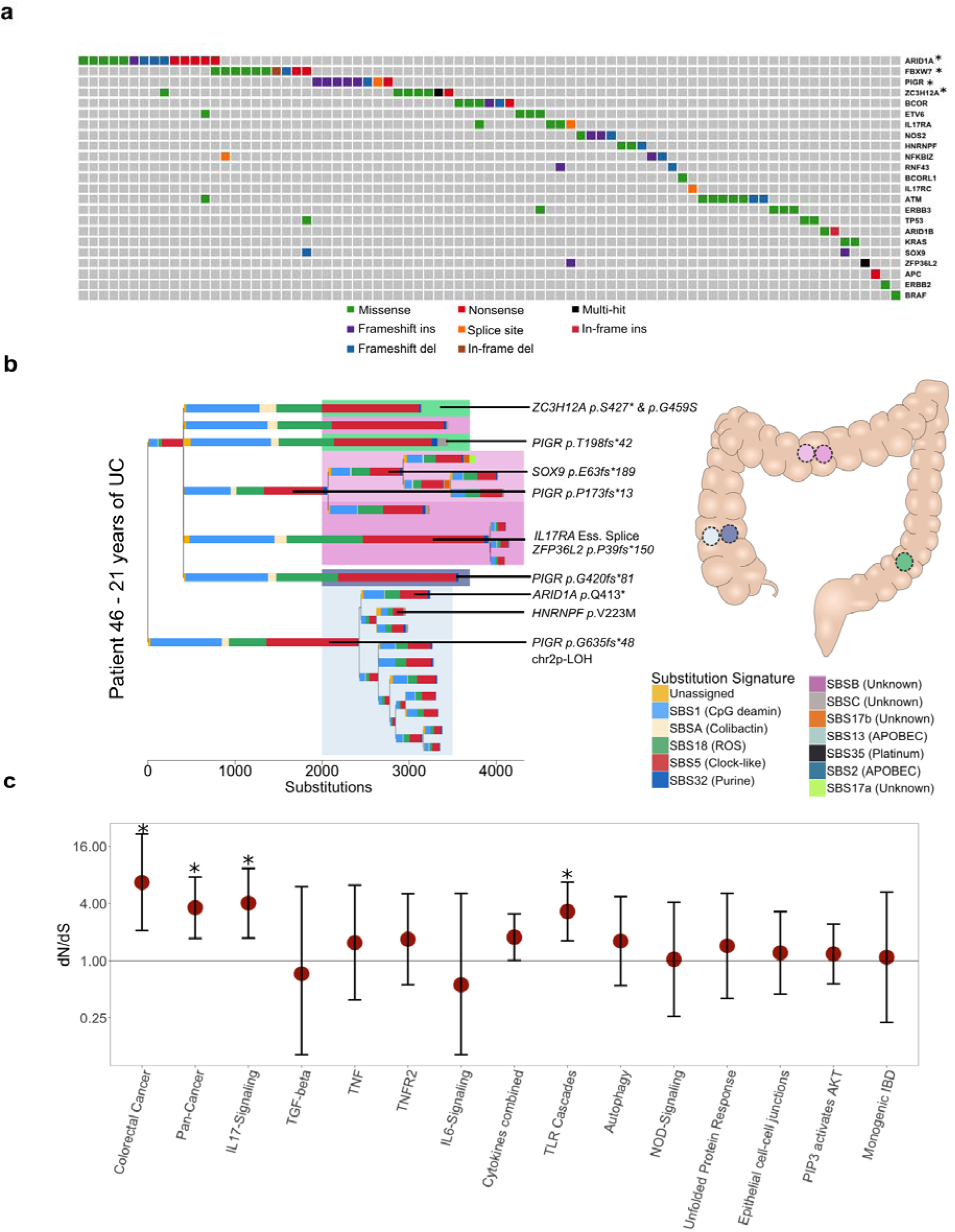
Driver mutations and positive selection in IBD. A) An oncoplot showing the distribution of potential driver mutations mapped to branches of phylogenetic trees. Each column represents a branch of a phylogenetic tree and a mutation may be found in multiple crypts if the branch precedes a clonal expansion. Branches without potential drivers are not shown for simplicity. *Genes significantly enriched in non-synonymous coding mutations. B) A phylogenetic tree of the crypts dissected from a 38 year old male suffering from UC for 21 years. Crypts are dissected from five biopsies from three previously inflamed sites of the colon. Crypts carrying distinct *PIGR* truncating mutations are found in four of the biopsies and in all three colonic sites. C) Pathway-level dN/dS ratios for truncating mutations in known cancer genes and cellular pathways important in IBD pathogenesis. Error bars represent 95% confidence intervals. *Significant enrichment of mutations after Benjamini Hochberg correction for multiple testing (q<0.05).

Recurrent mutations in *PIGR* and *ZC3H12A* are of particular interest since these have not been described in cancer but have roles in immunoregulation and reflect distinct mechanisms of positive selection in the IBD colon. *PIGR* encodes the poly-immunoglobulin (Ig) receptor, which transfers polymeric Igs produced by plasma cells in the mucosal wall across the epithelium to be secreted into the intestinal lumen (Johansen and Kaetzel, 2011). *Pigr* knock-out mice exhibit decreased epithelial barrier integrity and increased susceptibility to mucosal infections and penetration of commensal bacteria into tissues (Johansen *et al*., 1999). *ZC3H12A* encodes an RNAse, Regnase-1 (also known as *MCPIP1*). It is activated in response to TLR stimulation and degrades mRNA of many downstream immune signaling genes (Matsushita *et al*., 2009), including *PIGR* (Nakatsuka *et al*., 2018)*, NFKBIZ (Mino et al., 2015)* and members of the IL17 pathway (Garg *et al*., 2015). Four of the mutations in *ZC3H12A* occur in a DSGxxS motif which when phosphorylated marks the protein for ubiquitin-mediated degradation. Mutations of the corresponding residues in mice attenuate the phosphorylation (Iwasaki *et al*., 2011) and stabilize the protein so these are likely gain of function.

We next carried out a pathway-level dN/dS analysis, searching for enrichment of missense and truncating variants across 15 gene sets that were defined *a priori* because of their relevance in either colorectal carcinogenesis or IBD pathology (Figure 6C, Supplementary Tables 12, 13 and 14, Methods). We observed a 6.5-fold (1.8 - 23.6, 95% CI) enrichment of truncating mutations in genes associated with colorectal cancer (q=0.011) as well as a 1.9-fold (1.3-2.8, 95% CI) enrichment in genes significant in a pan-cancer analysis of selection (Priestley *et al*., 2019) (q=0.011). Interestingly, the pathway-level dNdS also revealed a 4.0-fold (1.7-9.4, 95% CI) enrichment of truncating mutations in the interleukin-17 (IL17) signaling pathway (q=0.011) and a 3.3-fold (1.6-6.7, 95% CI) enrichment in Toll-like receptor (TLR) cascades (q=0.011) with mutations from both UC and CD derived crypts contributing to the enrichment (Supplementary Figure 10).

## Discussion

We have used whole genome sequencing of individual colonic crypts to provide the most accurate characterization of the somatic mutation landscape of the IBD affected colon to date. Our results suggest that somatic mutagenesis of the mucosa is accelerated 2.4-fold in disease and that this increase is mostly driven by acceleration of common mutational processes which are associated with metabolic stress and are ubiquitous in IBD-unaffected colon. Metabolic stress also results in an increased burden of somatic structural variants, which nevertheless remain rare in the IBD-affected mucosa. Structural variants are common in colorectal cancers and thus rapid increase in structural variation may be a hallmark of neoplastic transition, in line with previous reports (Baker *et al*., 2018). Increase in structural variation from healthy tissue to non-neoplastic disease has also been observed in liver disease (Brunner *et al*., 2019).

Colitis-associated colorectal cancers commonly occur on the background of large clonal fields (Choi *et al*., 2017). In our sample of non-dysplastic tissue we find millimeter scale clonal expansions, although we note that for many inflamed regions only a single small biopsy is available which limits our ability to detect large clones. *TP53* and *KRAS* mutations are thought to be key events in clonal spread in the IBD mucosa but while we do observe a number of canonical cancer driver mutations in genes including *TP53* and *KRAS,* only *ARID1A* and *FBXW7* show significant evidence of positive selection.

While there is substantial overlap in the driver landscape of IBD and non-IBD colon, important differences also exist. Our findings of enrichment of mutations in *PIGR*, *ZC3H12A* and in the IL17 and TLR pathways suggest there are distinct selection mechanisms in the colitis-affected colon and that somatic mutations potentially play a causal role in the pathogenesis of IBD. While this work was under review, two studies of somatic mutations in UC patients from the Japanese population were published which confirm our findings of positive selection of mutations in *ARID1A, FBXW7, PIGR, ZC3H12A* and in the IL17-pathway (Kakiuchi *et al*., 2020; Nanki *et al*., 2020). Importantly, our study shows that the same selective pressures are operative in mucosal tissue in both ulcerative colitis and Crohńs disease.

The two papers also report mutations in additional genes including *NFKBIZ, IL17RA, TRAF3IP2 and NOS2*. We performed restricted-hypothesis testing of a set of 13 genes reported in these other two papers and replicate six at q<0.05 (Supplementary Table 15). Importantly, the enrichments of truncating mutations we observe in the IL17 and TLR pathways, which share many genes in common, are not driven by the genes discussed above because *PIGR, ZC3H12A, NFKBIZ* and *NOS2* are not part of these pathways (according to Reactome), and no mutations were found in *TRAF3IP2.* This suggests that additional positively selected genes related to IL17 and TLR signaling may be discovered in the IBD colon as sample size is increased. The difference in the number of *NFKBIZ* mutant crypts between the studies is noticeable. We detect only 3 truncating mutations in *NFKBIZ,* which is the most commonly mutated gene in Kakiuchi et al (Kakiuchi *et al*., 2020). This is reminiscent of our previous description of how selection of *NOTCH2* mutants in normal skin varies between individuals of European and South Asian ancestry (Martincorena *et al*., 2015). Together with our observation that distinct mutations in the same gene are often found in crypts from the same individual, this leads us to speculate that differences in local environment or a person’s genetic background affects the strength of selective advantage posed by somatic variants and studies with larger sample sizes may be able to detect those interactions.

In their study, Nanki et al show how IL17A may be cytotoxic to epithelial cells and argue that clones carrying IL17 pathway mutations are able to avert this cytotoxicity and thereby selectively expand in the inflamed environment (Nanki *et al*., 2020). This has implications for the direction of effect of these mutations on IBD pathogenesis since selective pressure would only be asserted following disease onset as Th17 cells infiltrate the tissue and secrete IL17A in the vicinity of the epithelium. However, it could also be hypothesized that these mutations play a causal role in the pathogenesis of IBD through an effect on dysbiosis. Indeed, the discovery by Nanki et at (Nanki *et al*., 2020) that *PIGR* mutations do not confer upon cells survival advantage in the presence of IL17A may add weight to this hypothesis. While *ZC3H12A* and *NFKBIZ* are involved in IL17 signaling, both are also induced downstream of TLRs (Yamamoto *et al*., 2004; Matsushita *et al*., 2009) where they regulate the transcriptional changes that follow TLR signaling. Disruption of the IL17 pathway itself may also play a causal role in the disease as intestinal epithelial-cell specific knock-out of components of the IL17 pathway in mice results in commensal dysbiosis through down-regulation of *Pigr* and other genes (Kumar *et al*., 2016). Thus, a positive feedback loop may be established, leading to ever greater spread of a pathogenic clone. It is worth noting that clinical trials of an anti-IL17A and anti-IL17RA antibodies for the treatment of Crohn’s disease have been carried out but either show no efficacy over placebo or worsen the disease (Hueber *et al*., 2012; Targan *et al*., 2016).

Our understanding of somatic evolution in normal tissues has improved greatly over the last few years but how and if somatic evolution contributes to the pathogenesis of complex traits other than cancer remains poorly understood. Clonal hematopoiesis has been associated with coronary heart disease (Jaiswal *et al*., 2014) and our work suggests that somatic evolution in the colonic mucosa may initiate, maintain or perpetuate IBD. Large scale analyses of cancers have started to reveal common themes of cancer evolution across tissues (Gerstung *et al*., 2020) and extending this work to other tissues exposed to chronic inflammation may similarly reveal patterns of remodeling of the selection landscapes associated with disease, but which need not drive neoplastic growth. Comparing the evolutionary forces in the IBD mucosa with those operating in psoriasis, celiac disease, asthma and other diseases affecting epithelial cells is an area of special interest.

## Supporting information

Supplementary Table 1 - Cohort characteristics.

Supplementary Table 2 - Crypt annotations, meta-data and mutation burden.

Supplementary Table 3 - Summary of the linear mixed effects models used to estimate the increase in mutation burden associated with IBD.

Supplementary Table 4a - All coding mutations detected in cases.

Supplementary Table 4a - All coding mutations detected in controls.

Supplementary Table 5 - All copy number changes that passed manual revision.

Supplementary Table 6 - All somatic retrotranspositions that passed manual revision.

Supplementary Table 7 - Canonical mutation hotspots associated with cancer development in the Cancer Genome Atlas.

Supplementary Table 8 - Putative driver mutations found in cases and controls.

Supplementary Table 9 - Driver association with mutational burden and mutational signatures.

Supplementary Table 10 - Full output of gene-level dNdS for cases and controls.

Supplementary Table 11 - Comparison of positive selection in healthy and the IBD-affected colon.

Supplementary Table 12 - Gene sets and pathways used for pathway dNdS.

Supplementary Table 13 - Pathway dNdS output for IBD and dNdS values for CD, UC and controls.

Supplementary Table 14 - Mutation counts for genes in the IL17 and TLR pathways. Counts are only from the IBD dataset.

Supplementary Table 15 - Restricted hypothesis testing of genes reported to be under positive selection in Kakiuchi et al (2020).

## Acknowledgements

This work was supported by the Wellcome Trust grant [206194]. We thank the staff of Wellcome Sanger Institute’s Sequencing, Sample Management and Informatics facilities for their contribution to the study. We thank Paul Scott for his help with the laser capture microdissection and Dr. Doug Winton for discussion on the design of the study and interpretation of its results. Finally, we give our dearest thanks to the participants of this study, who consented to having an invasive and sometimes painful colonoscopy procedure extended in order to donate samples for this study.

## Author contributions

SO, RM, MP, IM, TR, PJC and CAA designed the study. KA, CD, KS, MP and TR were involved in cohort ascertainment, phenotypic characterization and recruitment. SO, RM and YH embedded and sectioned biopsies and SO and RM also imaged, microdissected, and lysed colonic crypts with contributions from HLS and TB. MT reviewed biopsy histology. SO, TC, PR, TB, HLS and MS contributed to statistical analyses, mutation calling and filtering. HJ called and filtered somatic retrotranspositions. IM, PJC and CAA oversaw statistical analyses. SO, MRS, IM, TR, PJC and CAA interpreted the findings. SO wrote the first draft of the manuscript and all authors contributed to its final version. PJC and CAA supervised the study.

## Declaration of interests

CAA is a paid consultant for Genomics plc and Celgene. All other authors declare no competing interests.

## STAR Methods

### KEY RESOURCES TABLE

| REAGENT or RESOURCE | SOURCE | IDENTIFIER |
| --- | --- | --- |
| Software and Algorithms |  |  |
| Algorithms and software for calling somatic substitutions - CaVEMan. | Cancer Genome Project, Wellcome Trust Sanger Institute | http://cancerit.github.io/CaVEMan/ |
| Algorithms and software for calling somatic indels - PIndel. | Cancer Genome Project, Wellcome Trust Sanger Institute | http://cancerit.github.io/cgpPIndel/ |
| Algorithms and software for calling somatic retrotranspositions - TraFiC- | (Rodriguez-Martin *et al.*, 2020) | https://gitlab.com/mobilegenomesgroup/TraFiC |
| mem. |  |  |
| Algorithms and software for calling somatic copy number changes - BRASS. | Cancer Genome Project, Wellcome Trust Sanger Institute | https://github.com/cancerit/BRASS |
| Algorithms and software for calling somatic copy number changes - ASCAT. | Cancer Genome Project, Wellcome Trust Sanger Institute | https://github.com/cancerit/ascatNgs |
| Algorithms and software for constructing phylogenetic trees - MPBoot. | (Hoang *et al.*, 2018) | http://www.iqtree.org/mpboot/ |
| Algorithms and software for extracting mutational signatures - HDP. | (Roberts, 2018) | https://github.com/nicolaroberts/hdp |
| Algorithms and software for estimating dN/dS ratios. | (Martincorena *et al.*, 2017) | https://github.com/im3sanger/dndscv |
| Custom scripts documenting all other analyses. | This paper. | https://github.com/Solafsson/somaticIBD |
| Deposited Data |  |  |
| BAM files containing sequencing data from crypts isolated from IBD patients. | This paper. | European Genome-phenome Archive (EGA) - accession codes (TBC) |
| Images of microdissections and physical distances between crypts. | This paper. | Mendeley Data (TBC) |
| BAM files containing sequencing data from crypts isolated from controls. | (Lee-Six *et al.*, 2019) | European Genome-phenome Archive (EGA) - accession codes EGAD00001004192 and EGAD00001004193 |
| PCAWG mutational signatures. | (Alexandrov *et al.*, 2020) | https://cancer.sanger.ac.uk/cosmic/signatures |

### CONTACT FOR REAGENT AND RESOURCE SHARING

All requests for data and resources should be directed to the Lead Contact, Dr. Carl Anderson. All sequencing data will be made available via the European Genome Phenome Archive prior to publication (https://ega-archive.org/) and accession codes provided. Histology images and physical distances between dissected crypts will be uploaded to the Mendeley database.

### EXPERIMENTAL MODEL AND SUBJECT DETAILS

Colonic pinch-biopsies were donated by IBD patients undergoing regular surveillance of their disease at Addenbrooke’s hospital, Cambridge (Supplementary Table 1). All samples were obtained with informed consent of the donor and the study was approved by the National Health Service (NHS) Research Ethics Committee (Cambridge South, REC ID 17/EE/0338) and by the Wellcome Trust Sanger Institute Human Materials and Data Management Committee (approval number 17/113). We have complied with all relevant ethical regulations.

All donors are of white-European ancestry. The time between clinical diagnosis and date of biopsy was used to define the disease duration of a given individual. We added six months to this number for all patients because symptoms often precede diagnosis by several months and to avoid setting the disease duration to zero for patients who donated samples at the time of diagnosis. Time of purine treatment was estimated by consulting electronic health records from NHS databases. Biopsies were annotated as never, previously or actively inflamed using all available clinical data and NHS histopathology archives. The biopsy images (or an image of a second biopsy from the same site of the colon) were reviewed by a histopathologist. None of the patients had colorectal cancer, adenoma or dysplasia.

Biopsies from patients 1-26 were embedded in optimal cutting temperature (OCT) compound and sectioned, stained and fixed as previously described(Lee-Six *et al*., 2019). None of the samples were fixed in formalin. Subsequent biopsies were embedded in paraffin because this better preserved the morphology of the tissue. Biopsies were sectioned (10-20 μm PEN membrane slides (11600288, Leica) and stained with hematoxylin and eosin. Crypts were dissected using laser capture microdissection microscopy (LMD7000, Leica) and lyzed using ARCTURUS PicoPure DNA extraction kit (Applied Biosystems) according to the manufacturer’s instructions. DNA libraries were prepared as previously described(Lee-Six *et al*., 2019).

The control cohort was obtained from our previous publication on somatic mutations in the normal colon(Lee-Six *et al*., 2019). It consists of seven deceased organ donors, 31 individuals who underwent colonoscopy following a positive faecal occult blood test in a screening programme (16 of which were not found to have an adenoma or a carcinoma and 15 of which had colorectal carcinoma, although the biopsies used were distant from these lesions) and three paediatric patients who underwent colonoscopy to exclude IBD and who were found to have a histologically and macroscopically normal mucosa. We excluded one subject from the control cohort who had undergone chemotherapy and was a clear outlier in terms of mutation burden and showed an abnormal mutation profile.

## METHOD DETAILS

### Genome sequencing

Samples from patient 1 through 19 (Supplementary Tables 1 and 2) were whole genome sequenced on Illumina XTEN® machines as previously described(Lee-Six *et al*., 2019). Samples from other patients were whole genome sequenced on Illumina Htp NovaSeq 6000® machines using 150bp, paired end reads except for patients 60-62, which were whole exome sequenced on the same platform using the Human All Exon V5 bait set. Reads were aligned to the human reference genome (NCBI build37) using BWA-MEM.

### Mutation calling and filtering

#### Substitutions

Base substitution calling was carried out in three steps: Discovery, filtering and genotyping. Mutations were first called using the Cancer Variants through Expectation Maximisation (CaVEMan) algorithm(Jones *et al*., 2016). CaveMan uses a Bayesian classifier, incorporating base quality, read position, read orientation and more, to derive a posterior probability of all possible genotypes at every candidate site. Out of concern for field cancerization effect, patients 1 through 26, and patients from which only a few crypts were sequenced, were analysed using a matched normal sample dissected from non-epithelial tissue from one of the biopsies. As it became apparent that clones did not stretch between biopsies, we stopped sequencing non-epithelial tissue control samples from patients if crypts were dissected from multiple biopsies.

The substitution calls were next filtered, as previously(Lee-Six *et al*., 2019) described, to remove mapping artefacts, common single nucleotide polymorphisms and calls associated with the formation of cruciform DNA structures during library preparation. When matched normal samples were unavailable for the calling (see above), a large number of rarer germline variants remained post filtering. All sites where a somatic mutation was called in any crypt from a given patient were subsequently genotyped in all other samples from that patient by constructing read pileups and counting the number of mutant and wild-type reads. Only reads with a mapping quality of 30 or higher, and bases with a base quality of 30 or higher, were counted.

We next performed an exact binomial test to remove germline variants. True heterozygous germline variants should be present at a variant allele frequency (VAF) of 0.5 in all samples from an individual. Across all samples from a given individual, we aggregated variant and read counts at sites where a single nucleotide variant was called in at least one sample. We then used a one-sided exact binomial test to distinguish germline variants from somatic variants. The null hypothesis was that germline variants were drawn from a binomial distribution with a probability of success of 0.5, or 0.95 for the sex chromosomes in men. The alternative hypothesis was that these variants were drawn from distributions with a lower probability of success. The resulting p-values were corrected for multiple testing using the Benjamini-Hochberg method. A variant was classified as somatic if q <10^-3^, or q < 10^-2^ if fewer than five crypts had been dissected for the patient. For variants classified as somatic, we fitted a beta-binomial distribution to the number of variant supporting reads and total number of reads across crypts from the same patient. For every somatic variant, we determined the maximum likelihood overdispersion parameter (ρ) in a grid-based way (ranging the value of ρ from 10 to 10^-0.05^). A low overdispersion captures artefactual variants because they appear to be randomly distributed across samples and can be modelled as being drawn from a binomial distribution. In contrast, true somatic variants will be present at at a VAF close to 0.5 in some, but not in all crypt genomes, and are thus best represented by a beta-binomial with a high overdispersion. To distinguish artefacts from true variants, we used ρ= 0.1 as a threshold, below which variants were considered artefacts. The code for this filtering approach is an adaptation of the Shearwater variant caller(Gerstung, Papaemmanuil and Campbell, 2014). Finally, we filtered out variants that were supported by fewer than three reads or where the sequencing depth was less than five.

#### Indels

Short deletions and insertions were called using the Pindel algorithm(Ye *et al*., 2009). We applied the same restrictions on median VAF and read counts as for substitutions, and germline indel calls were filtered using the same binomial filters as described above.

#### Constructing phylogenic trees

We used the MPBoot software(Hoang *et al*., 2018) to create a phylogenic tree for each patient. MPBoot uses ultrafast bootstrap approximation to generate a maximum parsimony consensus tree. We assigned mutations to branches using a maximum likelihood approach, removing mutations which didn’t adhere to the tree structure (P<0.01, maximum likelihood estimation).

#### Structural variants

Copy number variants were called using the BRASS algorithm (https://github.com/cancerit/BRASS) as previously described(Lee-Six *et al*., 2019). Calls were filtered using AnnotateBRASS (https://github.com/MathijsSanders/AnnotateBRASS) as previously described(Moore *et al*., 2018). When a matched normal sample was not available for a patient, we used a clonally unrelated sample from the same individual to filter germline variants. All variants passing filters were manually reviewed in a genome browser. For discovery of deletions at fragile sites of the genome, we manually reviewed the three regions in all the genomes.

Somatic retrotranspostions were called using TraFic algorithm (Rodriguez-Martin *et al*., 2020). Somatic events supported by read clusters without exact breakpoints were also included. To further identify somatic transduction events, translocation calls (i.e., read clusters) related with known L1 germline sources (Rodriguez-Martin *et al*., 2020) from BRASS algorithm were examined. All somatic retrotransposition events were manually reviewed.

Chromosome aneuploidies and deletions or duplications affecting large areas of chromosomes or whole chromosome arms were called using the ASCAT algorithm(Van Loo *et al*., 2010; Raine *et al*., 2016)

### Mutation rate comparisons between IBD patients and controls

Any test for a difference in mutation burden between cohorts must take into account all factors, biological and technical, which correlate with disease and/or affect mutation calling sensitivity. For our comparison of IBD and normal, we fitted a linear mixed effects model taking the following factors into account:

1. Age is the most important predictor of mutation burden and the age distribution of the two cohorts is different. We include a fixed effect for age in our model to account for this.
2. Mutation burden differs for different sectors of the colon(Lee-Six *et al*., 2019). The IBD cohort is enriched with samples from the left side, as this is the area predominantly affected in UC patients. We include a fixed effect for location within the colon to account for this.
3. Observations are non-independent. We include in the model random effects for patient and for biopsy, with the random effect for biopsy nested within that for the patient.
4. Most embryonic mutations will be removed by our filters as germline so at birth the mutation count is near zero. We therefore do not include a random intercept in our model but constrain the intercept to zero. The biological interpretation of this is that there are no somatic mutations present at birth.
5. The between-patient variance is likely greater in the IBD cohort as patients vary in the duration, extent and severity of their disease. The within-patient variance is also likely greater in the IBD cohort as biopsies taken from different sites of the colon vary in their disease exposure, number and duration of flares etc. To model this, we construct a general positive-definite variance-covariance matrix for the random effects of patient and biopsy by cohort.
6. Any difference in the clonality of the colon between IBD patients and controls will affect the relative sensitivity to detect somatic mutations. To account for this, we adjusted the branch lengths of the phylogenic trees and used the adjusted mutation counts as the response variable in our models. The adjustment was carried out as follows. Mutations with low variant allele frequencies (VAFs) will be missed at low coverage. Therefore, for each crypt, we first fitted a truncated binomial distribution to the VAF distribution of the crypt to estimate the true underlying median VAF (this is different from 0.5 because recent mutations may not yet have been fixed in the stem cell niche, and because of contamination of lymphocytes and other cells from the lamina propria, which do not carry the same somatic mutations as the epithelial cells). We next simulated 100,000 mutation call attempts by drawing the coverage of each call from a Poisson distribution, with the lambda set as the median coverage of the sample, and multiplying that with the median VAF estimate from the truncated binomial. The resulting value represents the number of reads that carry the mutated allele. We calculated sensitivity for the sample, S_s_, as the fraction of draws that resulted in four or more mutant reads, which is the number required by CaVEMan to call a mutation. The sensitivity of a branch with n daughter crypts, S_b_, was then calculated as: S_b_ = 1 - (1-S_s1_)*...*(1-S_si_)*...*(1-S_sn_)

The adjusted mutation count is thus the observed mutation count divided by the sensitivity of the branch. In this way, the mutation count of clones formed of stand-alone crypts is augmented more than that of branches with multiple daughter crypts. Even after these steps, a small but significant effect of coverage remained and a fixed effect for coverage was included in the model.

We compared this model with one which includes disease duration as a fixed effect using a likelihood ratio test. The disease durations for never inflamed regions of the colons of IBD patients was set to zero.

As comparatively few CNVs and retrotranspositions are found in our dataset, we used Poisson regression within a generalized linear mixed effects framework to test for differences in structural variant number between cases and controls. We included the same random and fixed effects described above for base substitutions and compared models with and without disease duration using a likelihood ratio test. The above statistical tests are two-sided as are all statistical tests performed in this manuscript. Full details and outputs of all statistical models used in this work are available in an R-markdown file accompanying the submission.

#### Mutational signature extraction and analyses

Define a mutational signature as a discrete probability distribution over a set of categorical mutation classes (for example, 96 classes for single base substitutions - according to the identity of the pyrimidine-mutated base pair, and the base 5′ and 3′ to it, see (Alexandrov *et al*., 2013, 2020). We extracted mutational signatures using a hierarchical Dirichlet process (Roberts, 2018) (HDP, see the hdp R package https://github.com/nicolaroberts/hdp). This has the advantage of allowing simultaneous fitting to existing signatures and discovery of new signatures. We pooled the control and the IBD data and extracted signatures from both datasets as previously described for indels and single base substitutions separately(Lee-Six *et al*., 2019). We mapped mutations to branches of a phylogenic tree and treated each branch with more than 50 mutations as a sample. We used signatures reported in colorectal cancer as priors and also included as priors signature 32, which is attributed to azathioprine therapy(Alexandrov *et al*., 2020), and signature 35, attributed to platinum-based chemotherapy, as there are patients in our cohort with history of using these drugs. Using the PCAWG terminology (Alexandrov *et al*., 2020), the priors used were SBS1, SBS2, SBS3, SBS5, SBS13, SBS16, SBS17a, SBS17b, SBS18, SBS25, SBS28, SBS30, SBS32, SBS35,SBS37, SBS40, SBS41, SBS43, SBS45 and SBS49 for substitutions and ID1, ID2, ID3, ID4, ID5, ID6, ID7, ID8, ID10 and ID14 for indels. We used expectation maximization to deconvolute the HDP components into known PCAWG signatures. The cosine similarity between the HDP component corresponding to SBS1 was <0.95 and we used expectation maximization to break the component down into PCAWG signatures. We then reconstituted the components using only those PCAWG signatures that accounted for >10% of the mutations (this was done so as to avoid overfitting). This helped resolve the correlation between SBS1, SBS5 and SBS18. No other components had cosine similarity <0.95 with their corresponding signatures and other PCAWG signatures accounting for >10% of the mutations.

#### Selection analyses

To search for mutations under positive selection, we used the dNdScv method(Martincorena *et al*., 2017). We included never inflamed samples from the IBD cohort in the analysis as some uncertainty existed regarding the annotation of a handful of never-inflamed biopsies and we estimated that our analysis would suffer more from potential exclusion of drivers than from inclusion of more neutral mutations. We used the Benjamini-Hochberg method to correct for multiple testing.

To look for enrichment of mutations in pathways we defined 15 gene-sets (Supplementary Table 11) as follows. We included all genes found to be under selection in colorectal cancer(Priestley *et al*., 2019) and a list of genes significant in a pan-cancer analysis of solid tumours(Priestley *et al*., 2019). We also chose a set of cellular pathways known to be important in IBD pathogenesis and epithelial homeostasis. The Reactome database was used to define the pathways(Fabregat *et al*., 2018), see Supplementary table 11 for accession dates and Reactome IDs of pathways. We chose the cytokine pathways TNF-Signaling, TNFR2, IL6, TGFb and IL17 for testing. We also defined a combined list of cytokines which included all of the above as well as IFNg, IL10, IL20, IL23, IL28, and IL36. We also decided to test other pathways shown by us and others through genome-wide association studies to be important in IBD pathogenesis(Jostins *et al*., 2012; de Lange *et al*., 2017). These were Toll-like receptor cascades, NOD-signaling, autophagy, unfolded protein response and epithelial cell-cell junctions. We included the PIP3/AKT signaling pathways as it is downstream of many of the pathways defined above and we had discovered two deletions affecting this pathway before performing the analysis. Finally, we defined a list of genes known to cause early-onset, monogenic forms of IBD. Many of the genes defined in the literature affect myeloid cell development and cause severe immunodeficiencies(Uhlig, 2013; Uhlig *et al*., 2014). We restricted our analysis to the union of monogenic-IBD genes which either are specifically thought to affect epithelial cells or were members of any of the pathways above.

We extracted global dN/dS values for missense and truncating variants separately and used the Benjamini-Hochberg method to correct for multiple testing.

## Supplementary Figures and figure legends

**Supplementary Figure 1:**
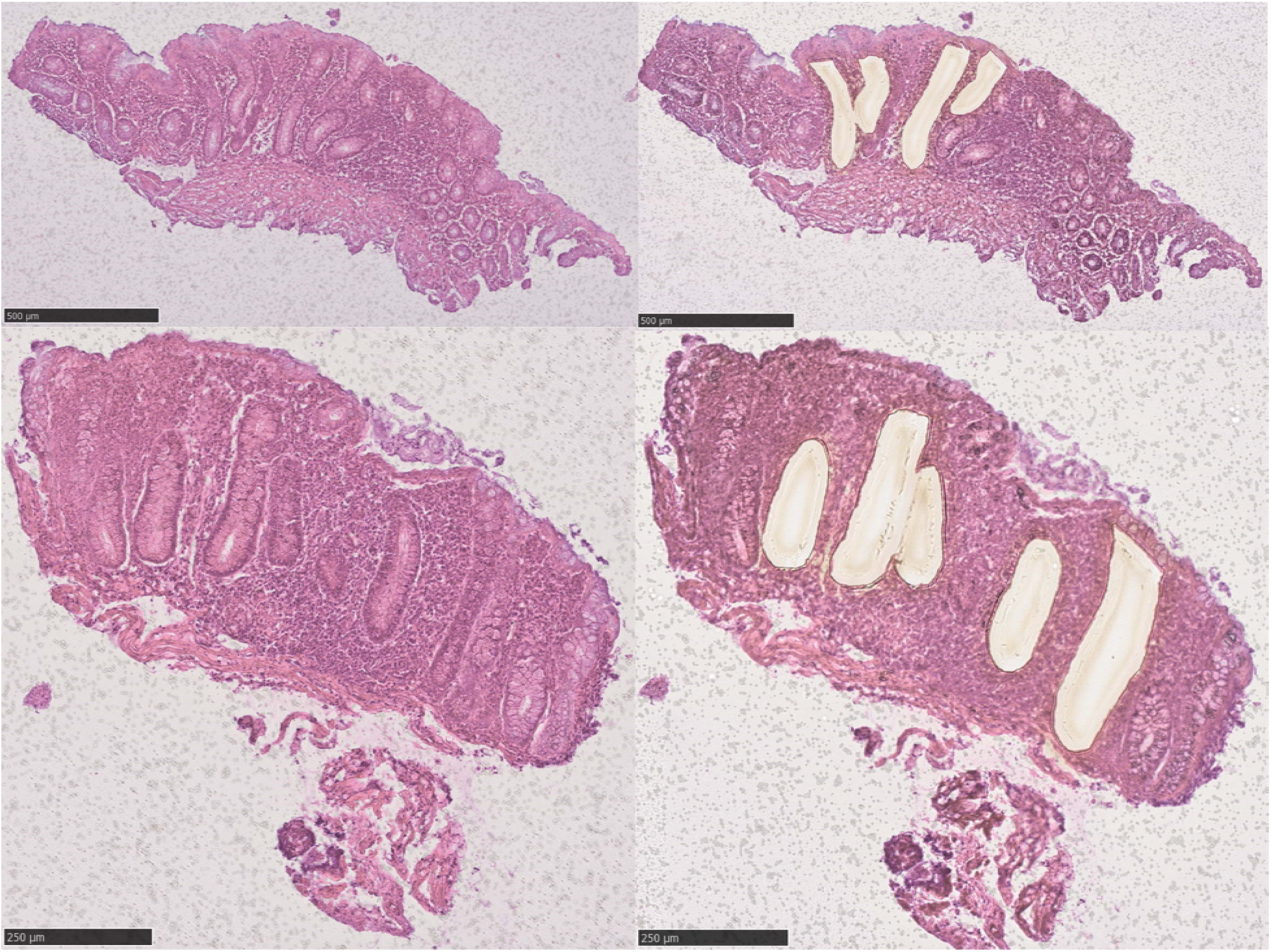
Laser capture microdissection of crypts from two biopsies. The left side shows representative images of sections of endoscopic biopsies from colonic tissue before crypt dissection. The right side shows the same sections after crypt dissection.

**Supplementary Figure 2:**
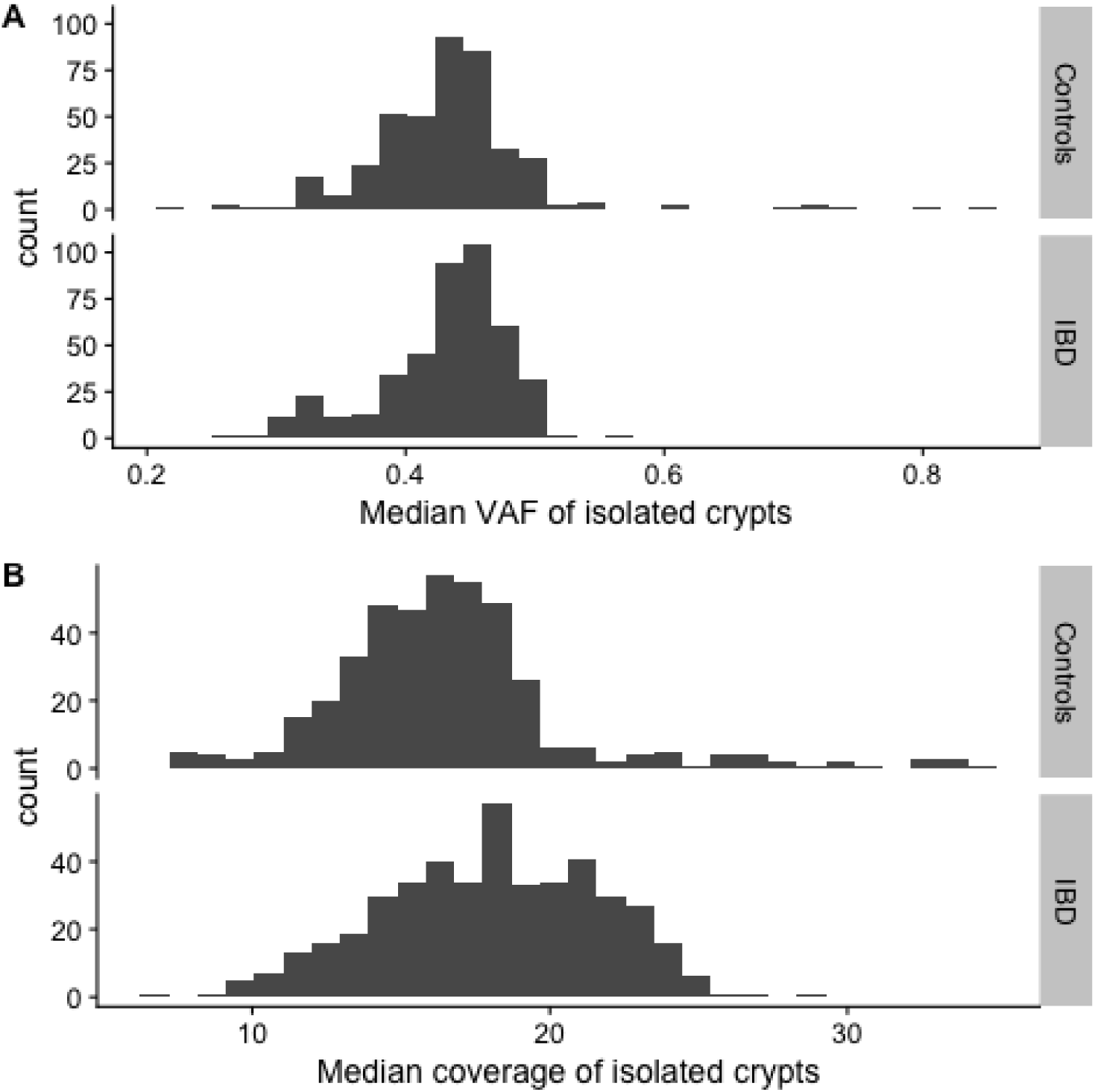
Median variant allele fraction (VAF) and coverage distributions for the IBD and control cohorts. The median-median VAF of the two cohorts is identical (0.44).

**Supplementary Figure 3:**
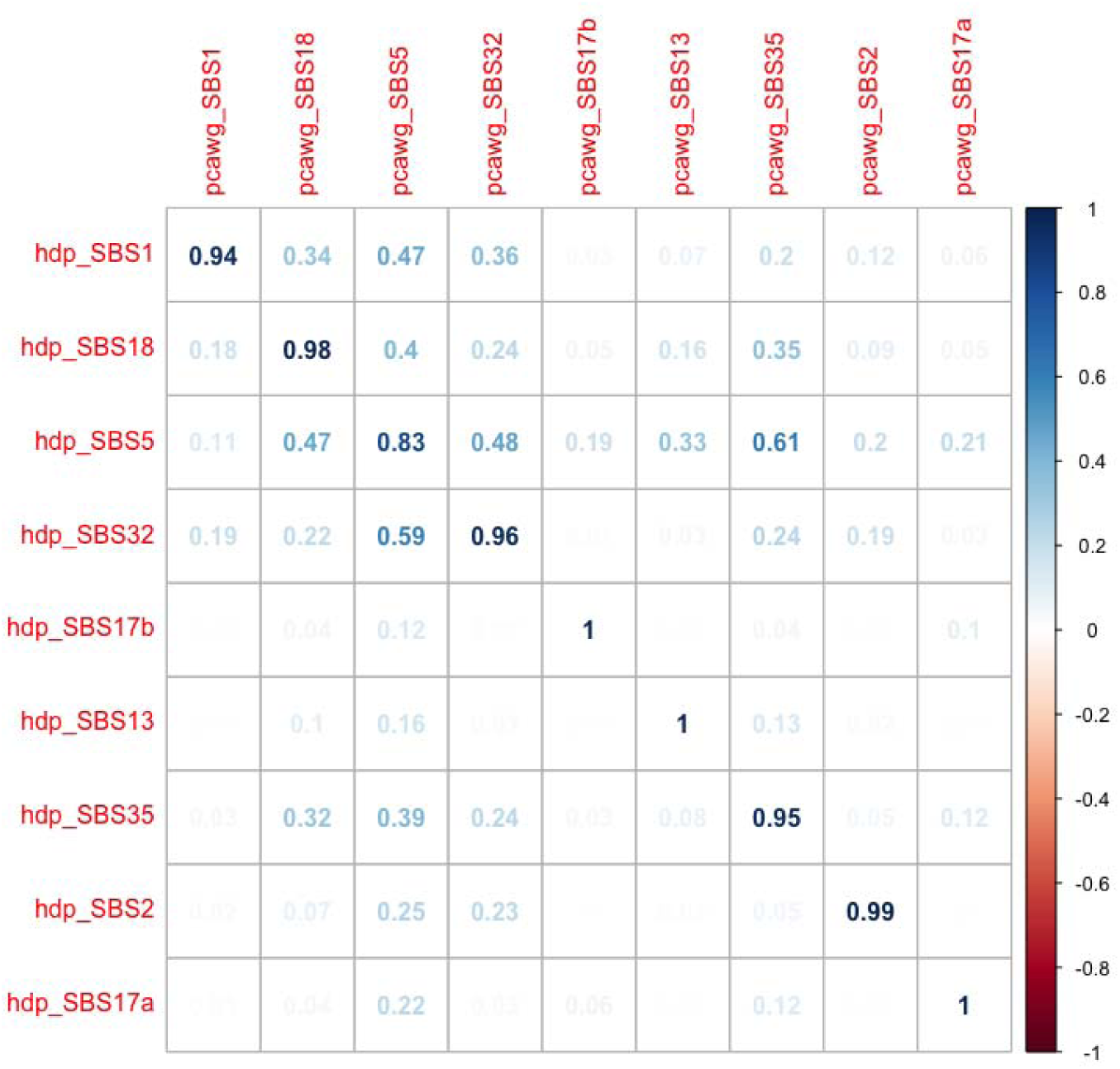
Cosine-similarities between single base substitution signatures extracted by hdp compared with published PCAWG signatures.

**Supplementary Figure 4:**
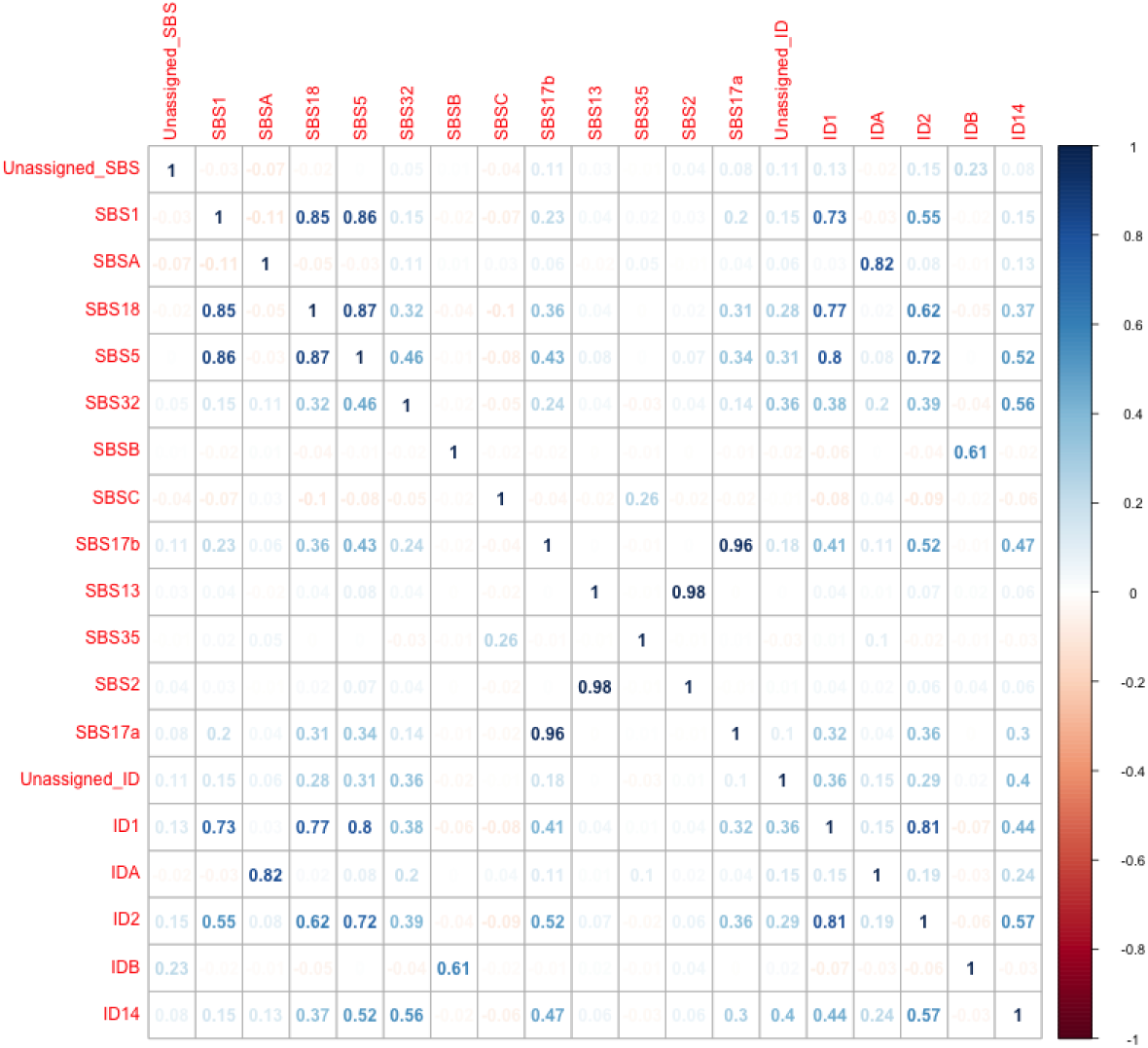
Correlation between identified signatures. SBSA is highly correlated with IDA and SBSB with IDB.

**Supplementary Figure 5:**
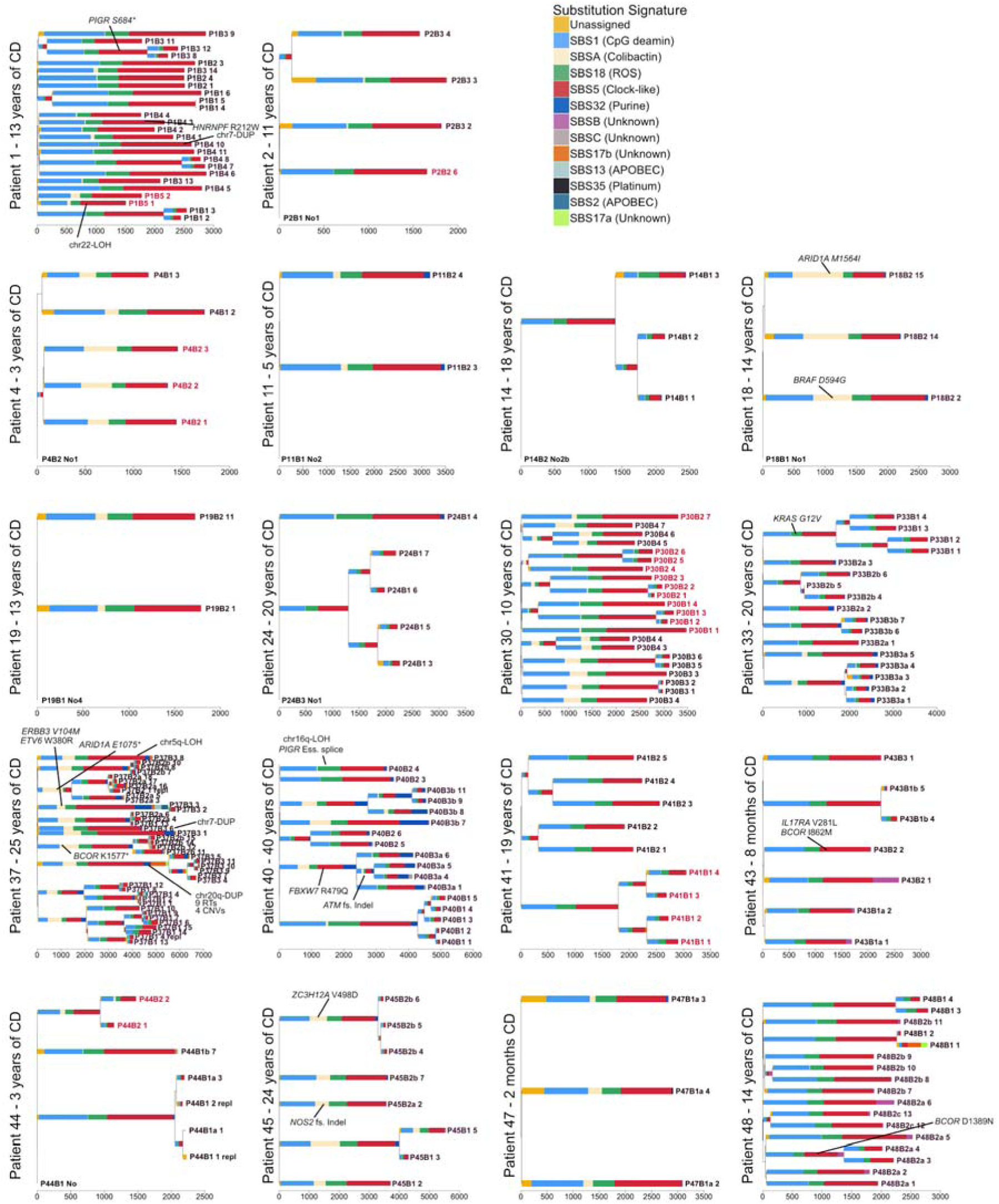
Phylogenetic trees for all Crohn’s disease patients. Mutational signatures are overlaid on the trees and likely driver mutations are mapped to the branch in which they occur. Crypts are labelled on the form PXBY_Z where PX is the patient number, BY the biopsy number (with a,b and c denoting biopsies taken a few millimeters apart from the same site) and Z is the crypt number. The colour of the labels indicates whether a crypt comes from an inflamed, previously inflamed or never inflamed site.

**Supplementary Figure 6:**
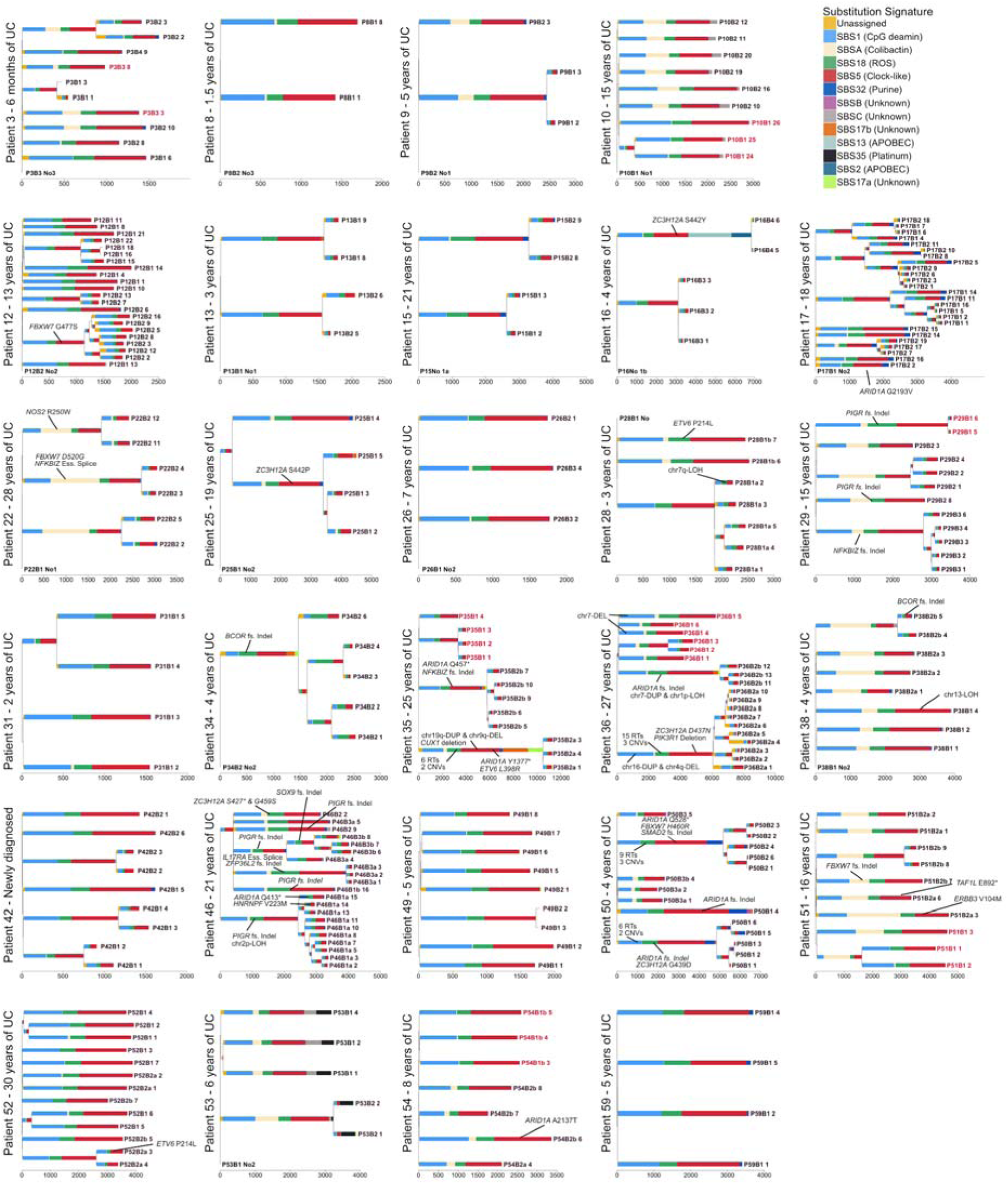
Phylogenetic trees for all ulcerative colitis patients. Mutational signatures are overlaid on the trees and likely driver mutations are mapped to the branch in which they occur. Crypts are labelled on the form PXBY_Z where PX is the patient number, BY the biopsy number (with a,b and c denoting biopsies taken a few millimeters apart from the same site) and Z is the crypt number. The colour of the labels indicates whether a crypt comes from an inflamed, previously inflamed or never inflamed site.

**Supplementary Figure 7:**
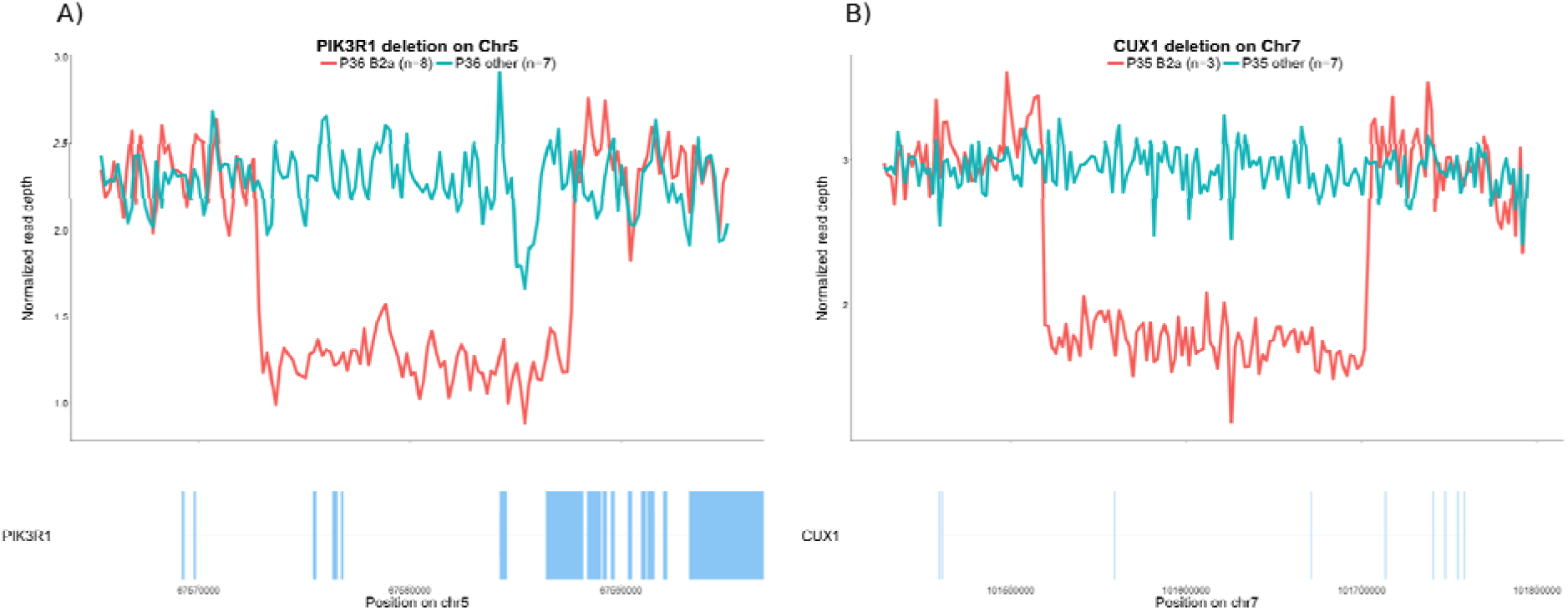
Structural variants of probable driver status. The figure compares normalized read depths of crypts called as carriers/non carriers. A) A deletion covering five exons of *PIK3R1* found to precede a clonal expansion in biopsy 2a of patient 36 (Figure 3 of the main text, middle panel, purple clone). B) A deletion covering three exons of *CUX1* and found to precede a clonal expansion in biopsy 2a of patient 35.

**Supplementary Figure 8:**
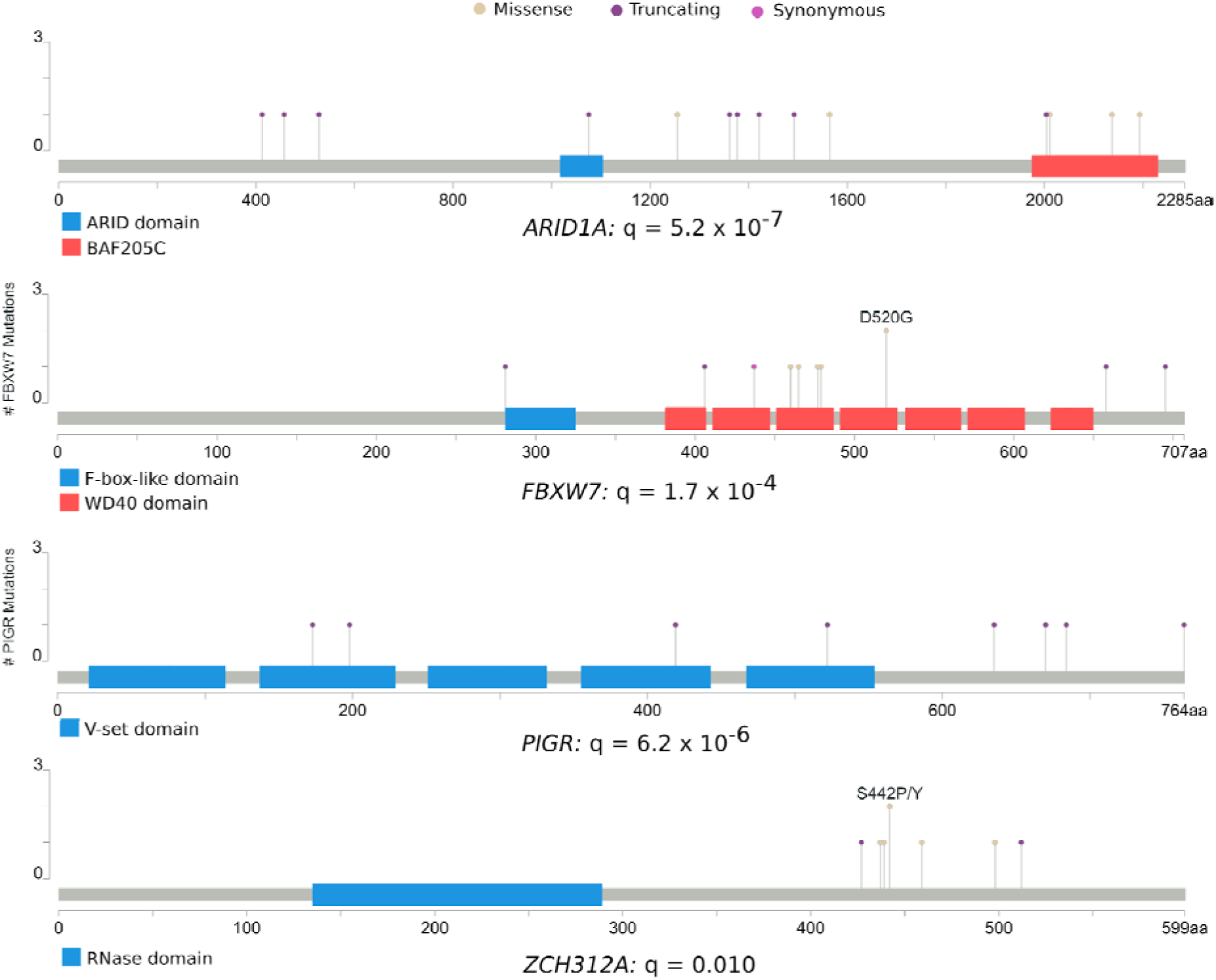
Mutations under positive selection. The plot shows the location of mutations found in genes that are enriched for non-synonymous coding mutations.

**Supplementary Figure 9:**
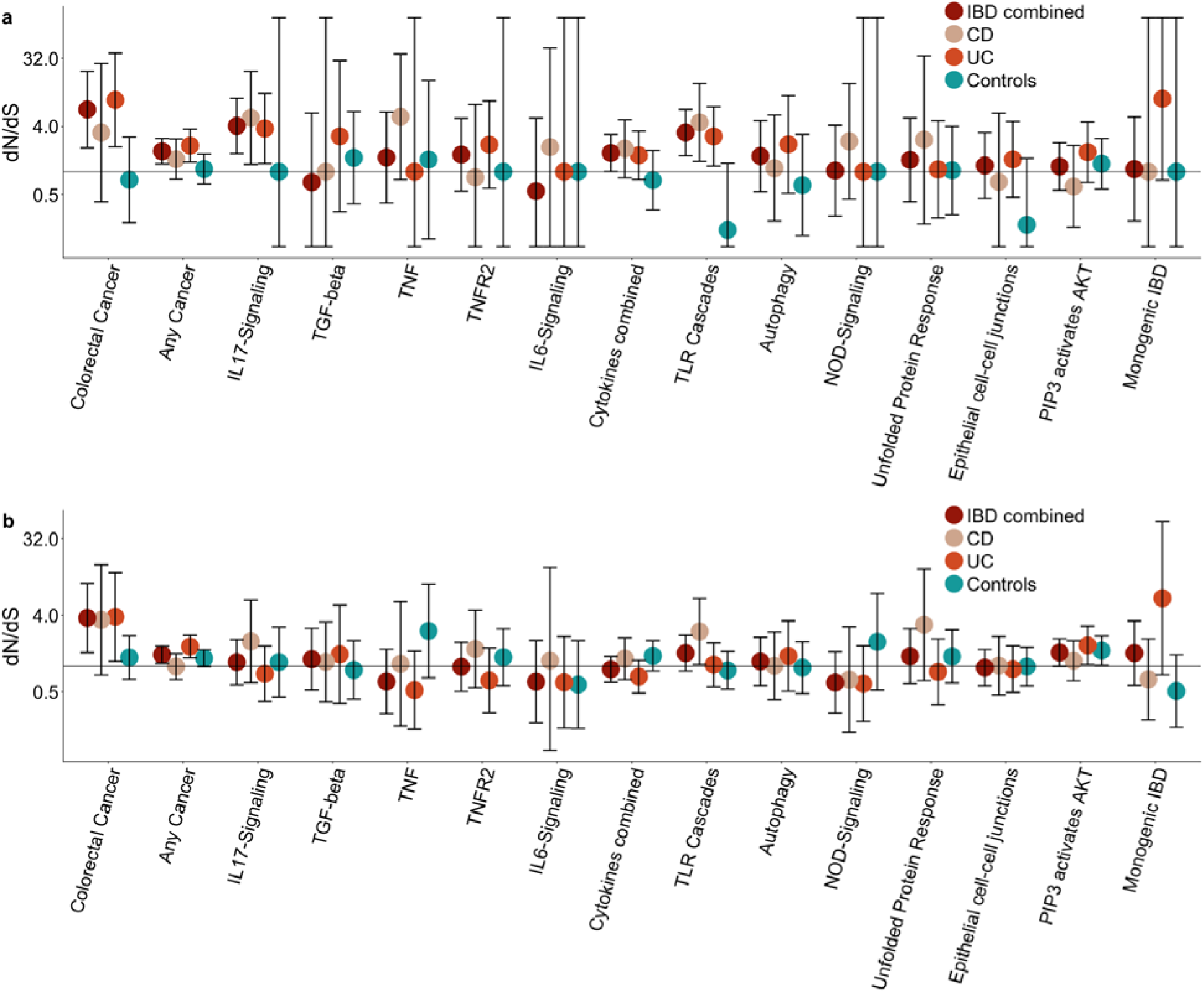
Pathway-level dN/dS ratios for mutations in known cancer genes and cellular pathways important in IBD pathogenesis. a) Pathway-level dN/dS for truncating mutations. Same as Figure 6c but also showing the ratios when analysis is restricted to CD or UC crypts. b) Pathway-level dN/dS for missense mutations.

